# Axolotl mandible regeneration occurs through mechanical gap closure and a shared regenerative program with the limb

**DOI:** 10.1101/2024.02.14.580108

**Authors:** Julia Kramer, Rita Aires, Sean D. Keeley, Tom Alexander Schröder, Günter Lauer, Tatiana Sandoval-Guzmán

**Affiliations:** Clinic of Oral and Maxillofacial Surgery, University Hospital Carl Gustav Carus Dresden, Technische Universität Dresden, Dresden, Germany; Department of Internal Medicine III, Center for Healthy Aging, University Hospital Carl Gustav Carus, Technische Universität Dresden, Dresden, Germany; Paul Langerhans Institute Dresden, Helmholtz Centre Munich, University Hospital Carl Gustav Carus, Technische Universität Dresden, Dresden, Germany

**Keywords:** axolotl, jaw, limb, regeneration, cartilage, gap minimization

## Abstract

The mandible plays an essential part in human life and, thus, defects in this structure can dramatically impair the quality of life in patients. Axolotls, unlike humans, are capable of regenerating their lower jaws; however, the underlying mechanisms and their similarity to those in limb regeneration are unknown. In this work, we used morphological, histological, and transcriptomic approaches to analyze the regeneration of lateral resection defects in the axolotl mandible. We found that this structure can regenerate all missing tissues in 90 days through gap minimization, blastema formation, and finally tissue growth, differentiation, and integration. Moreover, transcriptomic comparisons of regenerating mandibles and limbs showed that they share molecular phases of regeneration, that these similarities peak during blastema stages, and that mandible regeneration occurs at a slower pacing.

Altogether, our study demonstrates the existence of a shared regenerative program used in two different regenerating body structures with different embryonic origins in the axolotl, and contributes to our understanding of the minimum requirements for a successful regeneration in vertebrates, bringing us closer to understand similar lesions in human mandibles.

## Introduction

The jaws form the structural base of the mid and lower face and are essential for basic human activities like mastication and communication, as well as having an aesthetic component with an important role in social integration. Therefore, the loss of jaw tissue due to injuries or cancer can dramatically impair the quality of human life. Before the development of surgical techniques for mandibular replacement, resection of the anterior mandibular body resulted in the so-called “Andy Gump deformity”. This condition, based on the depiction of a historical cartoon character missing their chin and lip, resulted in severe functional limitations in eating and speaking (Lilly et al., 2021). Indeed, the symphyseal (distal-most), and parasymphyseal (immediately adjacent lateral) areas of the mandible, in particular, are especially important from a medical point of view, given their propensity to be affected by trauma or invasion of malignant tumors (Atilgan et al., 2010; Eichberger et al., 2022). If left unaddressed, lateral mandibular defects can cause issues like severe asymmetry of the lower face that intensifies when the mouth is opened and size discrepancies in the maxilla and mandible (Cohen and Schultz, 1985).

There are many procedures that can be used to help repair jaw injuries, such as preventing the ingrowth of unwanted soft tissue (membrane techniques), introducing missing cells or signals into the defect, and alloplastic replacement with metallic CAD/CAM-implants (Nauth et al., 2018; Reitemeier et al., 2016). In the latter case, new bone surrounding mandible replacement implants has been reported histomorphologically and radiologically in animals models and isolated cases of humans (Bräuer et al., 2023; Markwardt et al., 2014). Yet, the current gold standard for the treatment of critical-size segmental defects in the jaw is the grafting of suitable transplants from autologous donor regions (e.g., from the lower leg, hip, or shoulder blade) into the affected area (Disa and Cordeiro, 2000; Schemitsch, 2017). However, all these procedures are technically challenging, and the grafting of complex autogenous tissue transplants creates an extra injury at the donor site. Additionally, reconstructive surgery often provides just a compromise between aesthetics and function for the patient, and requires multiple treatments and long periods of hospitalization (Disa and Cordeiro, 2000; Kakarala et al., 2018).

In contrast to humans, certain vertebrate lineages, such as salamanders and teleost fish, possess the ability to completely reconstitute lost structures after amputation, including the lower jaw (Simon and Tanaka, 2013; Tanaka, 2016). Therefore, a systematic examination of jaw regeneration in highly regenerative model organisms is a promising approach to discover innovative therapeutic strategies.

Research in the newt *Notophthalmus viridescens* has helped to illustrate the general principles governing regeneration of the upper (maxilla) and lower (mandible) jaws. In newts, both adult and larval mandibles can regenerate following either full transections of the entire distal region (Ghosh et al., 1994; Goss and Stagg, 1958), or partial parasymphyseal resections (Graver, 1973; Graver, 1974; Graver, 1978), with both amputation types triggering a similar sequence of events. In adult newts, regeneration begins with the retraction of the severed mandible muscles and the healing of the wound. This is followed by the formation and thickening of the wound epidermis, the appearance of a blastema, cartilage differentiation, and, in the case of transection amputations, the joining of the newly regenerated Meckel’s cartilage in the midline’s medial symphysis. Regeneration of the bony components starts via formation of the dentary bone on the lateral sides of the regenerated cartilage and the re-formation of teeth. After approximately 20 weeks, the regenerated mandible resembles the unamputated jaws in both shape and size. Interestingly, larvae regenerate faster than adult newts and are able to reconstitute the prearticular bone and the lingual side of the dentary bone, whereas these structures never regenerate in the adult (Ghosh et al., 1994; Goss and Stagg, 1958).

Taking advantage of the recent development of many genetic resources and tools (Echeverri et al., 2022), we used the axolotl (*Ambystoma mexicanum*) to investigate jaw regeneration, especially in the context of limb regeneration. The limb stands as the reference example of blastema-dependent regeneration of a complex, multi-tissue structure, having been extensively studied throughout the years (reviewed in Aires et al., 2020; Payzin-Dogru and Whited, 2018). This structure also shares morphological features with newt jaw regeneration. Namely, limb regeneration begins with wound closure by a specialized wound epithelium (WE) that will later proliferate into a multilayered Apical Epidermal Cap (AEC), followed by blastema formation, tissue patterning and differentiation, in which cartilage and bone are formed, and, finally, a growth phase into a fully regenerated limb. As the molecular underpinnings of each of these processes are relatively well-known in the axolotl, limb regeneration is, then, the ideal blueprint for comparative studies between different regenerative structures in the same animal. Moreover, studying axolotl mandible and limb regeneration enables the comparison of regenerative processes of two structures with different embryonic origins, as the jaws derive from migrating neural crest cells, whereas limbs develop from the embryo’s lateral plate mesoderm (Gilbert, 1997).

However, contrary to limb regeneration, studies of mandible regeneration specifically in the axolotl are scarce, being so far mostly focused on tooth regeneration (Makanae et al., 2020) or in the analysis of punch biopsy lesions (Charbonneau et al., 2016; Charbonneau et al., 2021). In this work, we focused on an injury type that resembles defects in human patients that generally arise in the context of tumor surgery as, in cases like squamous cell carcinomas, it is sometimes necessary to perform large resections that often result in extensive full-thickness segmental defects. Specifically, we used a full-thickness lateral resection model that included skin, muscle, oral mucosa, and the medial symphysis with its laterally adjacent region. By analyzing the morphological, histological, and transcriptomic aspects of symphyseal and parasymphyseal mandible regeneration in the axolotl, we were able to examine the regenerative potential of both distal and proximal skeletal stumps within a segmental defect, as well as overall tissue integration. We found that the mandible is able to regenerate most missing tissues through two phases of mechanical gap minimization, formation of a blastema, and tissue growth, differentiation, and integration into the pre-existing structures. Moreover, gene expression profile comparisons of regenerating lower jaws with regenerating limbs showed that they undergo similar molecular phases of regeneration, particularly during blastemal stages. Yet, we found that regeneration in the lower jaw seems to progress at a slower pace when compared to the limb, and that this pacing can even differ between the two skeletal stumps of the regenerating mandible. Overall, our study uncovers the molecular details involved in the regeneration of resection defects in the axolotl mandible and highlights mechanistic differences and similarities in the regenerative processes of different body structures with different embryonic origins, which point at at the existence of a shared regenerative program in the axolotl.

## Results

### Repair of lateral resection defects in the axolotl mandible involves two phases of gap minimization

In order to study full-thickness lateral defects in the symphyseal and parasymphyseal regions in the axolotl mandible, we resected a 5 mm fragment of the right hemi-mandible. The amputation began in the medial symphysis and comprised Meckel’s cartilage, surrounding bone, epidermis, loose connective tissue, muscle, and gingiva. This resulted in a gap bordered by proximal and distal bony stumps, the latter still containing the remnants of the medial symphysis of the contralateral side (Fig. 1A). This defect amounted, on average, to approximately 43.3% of the perimeter of the hemi-mandible, or 20.7% when considering the full mandible perimeter (Fig. 1B). We then examined the gross morphological progression of repair for 90 days (Fig.1, Fig. S1).

**Fig.1.**
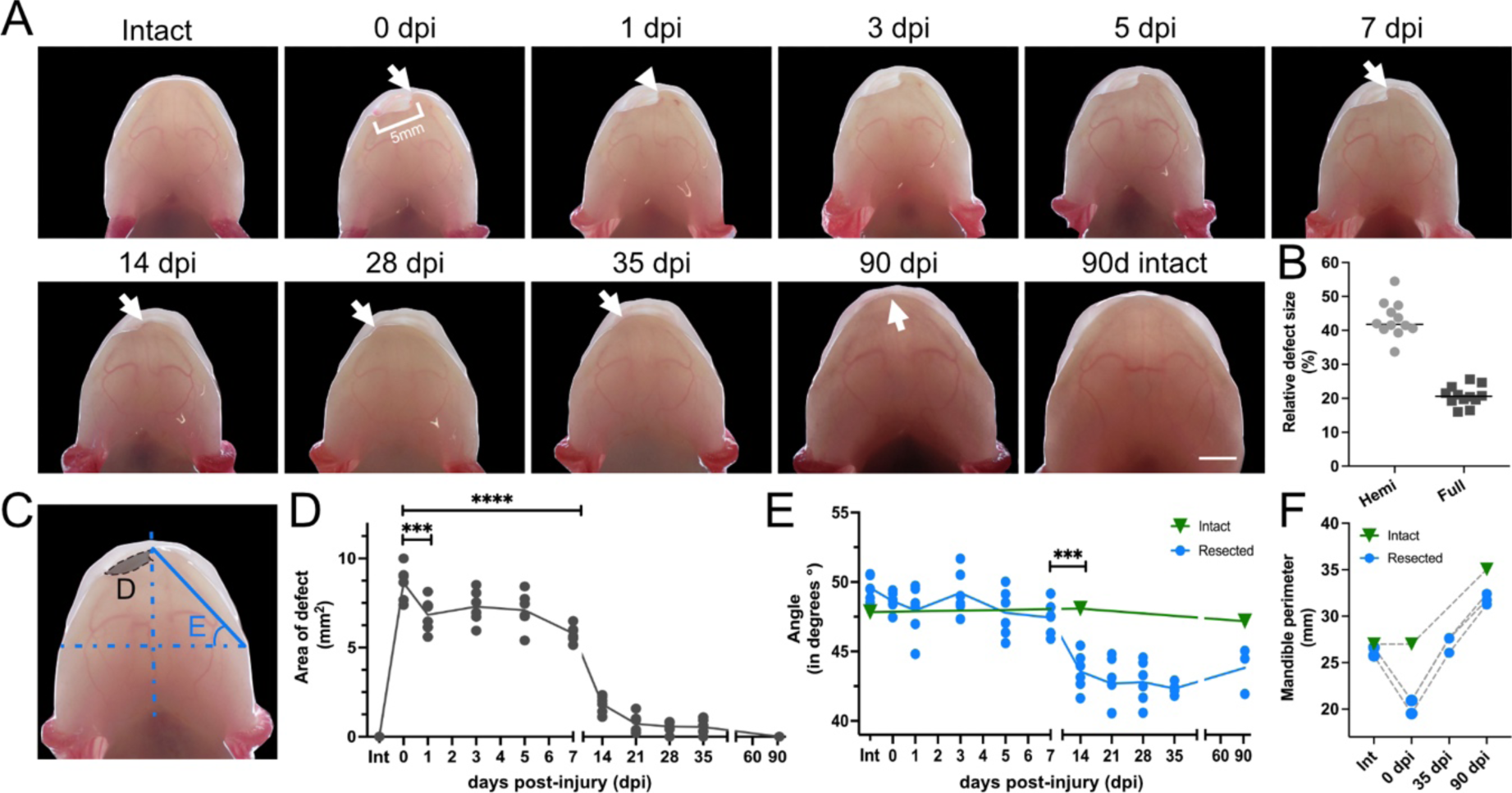
Repair of lateral resection defects in the axolotl mandible involves two phases of gap minimization. A. Time course of mandible regeneration after a full-thickness lateral resection. The same animal is shown immediately prior to injury (Intact), immediately following injury (0 dpi), and at 5-, 14-, 35-, and 90 dpi. Arrow and arrowheads indicate the distal stump of the injury. Scale bar: 5mm. **B.** Quantification of defect size relative to the perimeter of the hemi-mandible (Hemi) or the full mandible (Full) at 0 dpi. **C.** Schematic representation of defect area and distal stump angle quantifications. **D.** Quantification of defect area. Statistical significance was assessed via one-way ANOVA, ***=p<0.001, ****=p<0.0001. **E.** Angle of dislocation of the distal stump in resected (blue) and medial symphysis in intact (green) animals. Statistical significance between 7- and 14 dpi was assessed via one-way ANOVA, ***=p<0.001. **F.** Quantification of full mandible perimeter of intact and resected animals immediately before (Int) and post resection (0 dpi), and at 35- and 90 dpi.

Within the first day post-injury (dpi), we observed that the wound had stopped bleeding and that the overall defect region showed some signs of tissue contraction, as evidenced by a reduced concavity of the associated soft connective tissue and an apparent sharpening of the distal stump of the injury (Fig. 1A, white arrowhead; Fig. S1). The defect became less prominent at 14 dpi, was mostly gone at 35 dpi, and, by 90 dpi, the jaw seemed to have nearly regained its original shape (Fig. 1A). Quantification of the defect area (Fig. 1C) supported these initial observations, revealing that a significant defect minimization had already occurred within 1 dpi (Fig. 1D). The area of the defect then remained relatively unchanged until 5 dpi, but, by 7 dpi, had decreased again. By 14 dpi, the injury size was dramatically reduced and, by 35 dpi, had disappeared entirely (Fig. 1A, 1D, Fig. S1). However, the mandible had still not attained a proper shape in resected animals at this time point since, contrary to the intact controls, the lower dental arch appeared noticeably shorter (i.e., in a retrognathic mandibular position), resulting in a clear overbite of the upper jaw (Fig. 1A). We hypothesized that this shortening of the lower jaw could be due to movement of one or both mandibular fragments. To investigate this, we quantified the angle of displacement of the distal stump (Fig. 1C, 1E), which confirmed that this structure had moved laterally towards the defect, reaching a minimum angle (i.e., maximum displacement) at 35 dpi (Fig. 1A, white arrows; Fig. 1E). Remarkably, the angle of displacement increased afterwards, which suggested that the distal edge had moved towards the midline by 90 dpi and that the injured mandible was progressively regaining a normal shape. Indeed, by this time point, resected lower jaws seemed to have recovered their appropriate, orthognathic position in relation to their upper jaws (Fig. 1A). We confirmed this by measuring the perimeter of resected and intact mandibles, which showed that the initial mandible perimeter had been restored at 35 dpi, and was only 9.2% smaller when compared to the intact control at 90 dpi (Fig. 1F).

As such, our data demonstrates that, unlike humans but similar to other salamanders, the axolotl is capable of repairing lower jaw defects after a full-thickness lateral resection. We also found that their repair involved mechanical gap minimization in two main phases: the first occurring within the first day after injury and involving mainly tissue contraction; and the second starting at 7 dpi and involving tissue growth and displacement of the distal stump. Finally, we show that resected mandibles essentially recover their intact shape and size after 3 months. This strongly hints at a re-formation of most removed tissues by 90 dpi, especially cartilage and bone.

### Lateral defects in the mandible regenerate through tissue growth, differentiation, and integration of bone, cartilage and muscle

To investigate mandible regeneration in more detail, we performed histological stainings in intact (Fig. 2A, 2H) and resected animals at 5-, 14-, 35- and 90 dpi (Fig. 2B-G). The intact lower jaw of the axolotl consists of two paired cartilaginous rods known as the Meckel’s cartilage that extend through the entire length of the jaw in each side of the mandible and join at their distal ends by a median symphysis, in which the muscles of the floor of the mouth attach (Fig. 2A, A’; Fig. 2H, H’, gray arrowhead).

**Fig.2.**
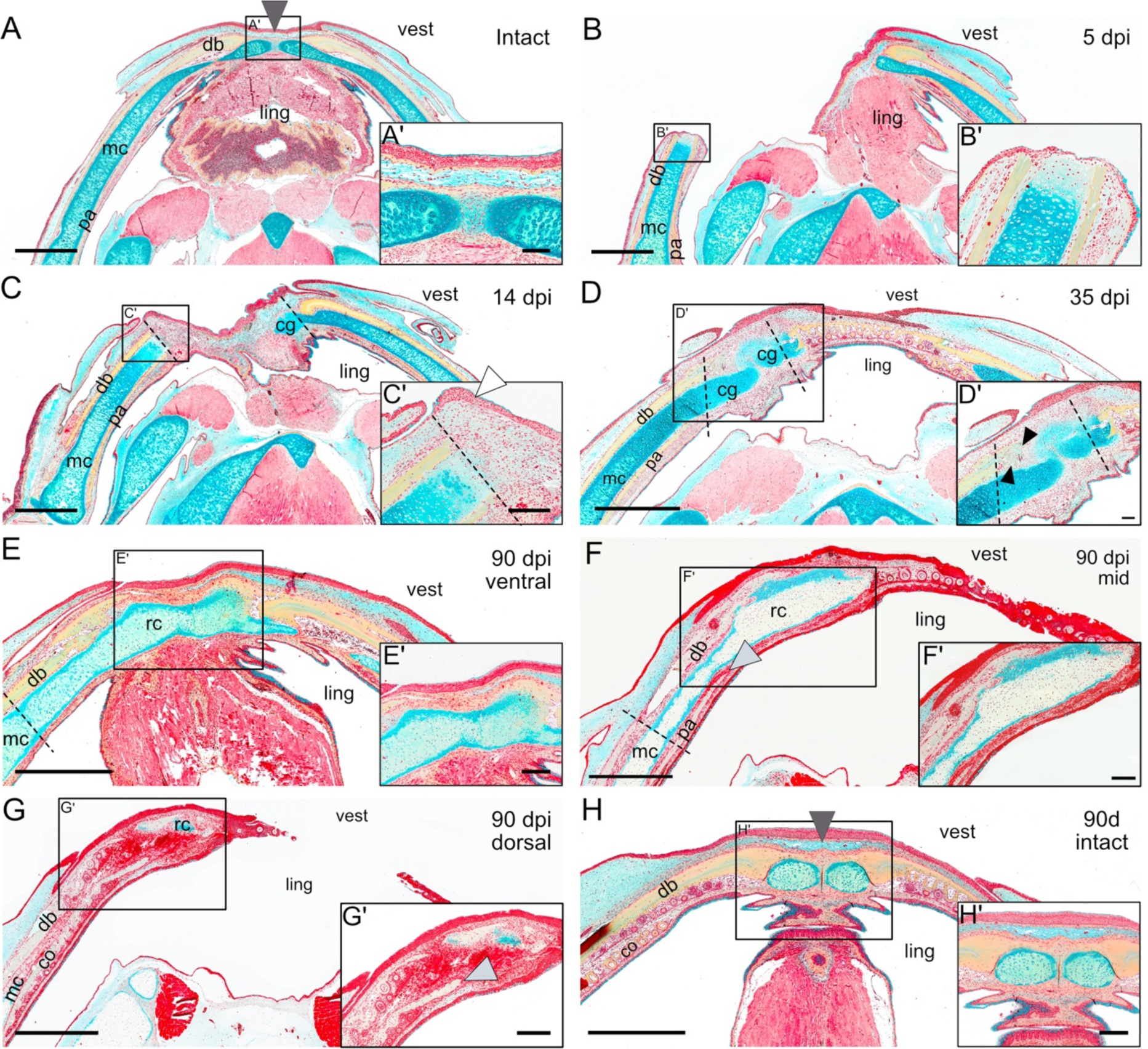
Lateral defects in the mandible regenerate through tissue growth, differentiation, and tissue integration of bone, cartilage and muscle. A-H. Movat’s pentachrome staining of longitudinal sections of intact mandibles at the beginning (A) and end (H) of the experiment, and resected mandibles at 5- (B), 14- (C), 35- (D), and 90 dpi (E-G). For 90 dpi, three sections corresponding to ventral (E), mid (F), and dorsal (G) levels are shown. Dark gray arrowheads in A and H indicate the medial symphysis in intact animals; white arrowhead in C’ shows a multilayered WE; black arrowheads in D’ point at intramembranous ossifications of the dentary bone; light gray arrowheads in F and G’ indicate the regenerated prearticular and coronoid bones, respectively. Dotted lines represent the approximate site of resection. mc: Meckel’s cartilage; pa: prearticular; co: coronoid; db: dentary; vest: vestibular side; ling: lingual side; cg: cartilaginous growth; rc: regenerated cartilage. Scale bar in A-H: 2 mm. Scale bar in insets: 500 µm.

Each Meckel’s cartilage rod is enclosed by several bones, including the dentary bone on its ventral and vestibular (or lateral) sides, the prearticular bone in the lingual side, and the coronoid bone resting above each prearticular bone. Teeth develop from both the dentary bone and the coronoid bone (Atkins et al., 2020; Makanae et al., 2020). This anatomy remains constant as the axolotl matures, unlike in the newt where the coronoid bone is lost during metamorphosis (Davit-Béal et al., 2007).

At 5 dpi, the hard tissue stump on the proximal side was covered by a thin multilayered epithelium (Fig. 2B, B’), under which the underlying cartilage showed changes in the extracellular matrix (ECM) and enlarged nuclei, both typical of histolysis and cell proliferation, respectively. Histolysis also occurred in the medial stump, as evidenced in one instance where the resection failed to go through the medial symphysis, instead going through the distal-most part of the same hemi-mandible (Fig. S2A, A’).

At 14 dpi, the gap caused by the resection was noticeably smaller, which was consistent with our macroscopic observations (Fig. 2C). Importantly, we found an accumulation of seemingly undifferentiated mesenchymal cells resembling a blastema sitting atop the injured stumps on both sides of the defect and spanning the full length of the gap (Fig. 2C, C’; Fig. S2B). These were entirely covered by a multilayered epithelium that was morphologically highly reminiscent of the AEC formed during limb regeneration, including the lack of a basement membrane (Neufeld and Aulthouse, 1986) (Fig. 2C, white arrowhead in C’; Fig. S2B, white arrowhead in B’ and B’’). Remarkably, resected proximal and distal stumps appeared to be in different regeneration stages. In the distal stump, a new cartilage growth that was continuous with the intact Meckel’s cartilage had already formed (Fig. 2C). This structure not only extended distally past the amputation plane into the defect, but also grew dorsally to surround the upward bend of the preexisting contralateral Meckel’s cartilage (Fig. S2B, B’’). This contrasted with the proximal stump, in which no cartilage condensations were yet detected. However, we did notice glycosaminoglycan filaments distal to its amputation plane, likely the first signs of cartilaginous extracellular matrix deposition (Fig. 2C, C’).

At 35 dpi, the gap caused by the resection was fully closed (Fig. 2D, Fig. S2C). The displacement of the distal stump towards the defect side was particularly evident, especially when compared to anatomical landmarks for the original lower jaw midline, such as the median raphe of the paired oral floor and the tongue muscles. The closing of the gap was accompanied by new cartilaginous growth from both sides of the defect, forming a continuous brace that traversed the length of the defect and integrated with the existing Meckel’s cartilage (Fig. 2D, D’; Fig. S2C, C’). The connection of these cartilage growths to each other, however, was imperfect, as they merged in a disorganized manner (Fig. 2D’). Interestingly, we found osteoid trabeculae with several cells trapped within the collagen deposits originating from the proximal stump (Fig. 2D’, black arrowheads; Fig. S2C’), indicating that bone reformation had already started to occur by intramembranous ossification.

Finally, by 90 dpi, the cartilage regenerate now robustly bridged the defect, having fully integrated into the previously existing Meckel’s cartilage in both sides of the defect and becoming the insertion site of the regenerated ventral mandibular muscles (Fig. 2E). Moreover, the formerly observed displacement of the midline towards the defect side was partially compensated (Fig. 2E, F; Fig. S2D). Additionally, the regenerated cartilage was thicker than the original Meckel’s cartilage of the injured region, and displayed some pockets of cartilaginous tissue surrounded by bony matrix (Fig. S2D, D’). We also noticed that, even though the newly regenerated dentary bone did not cover the entirety of the cartilage regenerate, it contained new teeth (Fig. 2G, G’). Similarly, we likewise saw that the prearticular and coronoid bones had extended past the amputation level (Fig. 2F, F’, Fig. 2G, G’, gray arrowheads) and that the latter contained both tooth buds and mature teeth, indicating that these structures had regenerated by 90 dpi. Finally, we interestingly did not see the reformation of the medial symphysis (Fig. 2E, Fig. 2F), with the regenerated cartilage brace instead connecting directly to the Meckel’s cartilage and to the dentary bone of the contralateral hemi-mandible.

Taken together, our results indicate that after a full-thickness lateral resection, the axolotl lower jaw can fully regenerate all missing tissues via the formation of a mesenchymal blastema-like structure, tissue growth, and tissue differentiation. Remarkably, the regenerate was also able to fully integrate into the previously existing mature structures on both sides of the defect, in such a way that the mandible shape was nearly restored after 90 days.

### Transcriptomic analysis of mandible regeneration after lateral resection uncovers significant similarities to limb regeneration

Next, we sought to get a better molecular understanding of the regeneration process in our mandible resection model by exploring its transcriptomic profile. For that, we performed bulk RNA-Seq in intact and regenerating mandible tissues in early (5 dpi), middle (14 dpi), and late (35 dpi) regeneration stages. Genes that were either lowly expressed or did not display any substantial variation over time were filtered out (see Materials and Methods). To visualize gene expression dynamics, the remaining genes were subjected to a *k*-means clustering algorithm, which subset them into 15 clusters based on their expression over time (Fig. S3, Table S1). Finally, genes that did not fit well into any of these clusters were removed due to concerns over their expression levels possibly arising from artificial or technical issues, ultimately leaving us with a list of 2,134 genes of interest. From the clusters containing these remaining genes, 10 of them fell into 4 broader categories of temporal expression patterns: clusters containing genes whose expression peaked at 5 dpi (*5 dpi Peak*), 14 dpi (*14 dpi Peak*), 35 dpi (*35 dpi Peak*), and that steadily rose over time (*General Rise*) (Fig. 3A, Fig. S4, Table S2).

**Fig.3.**
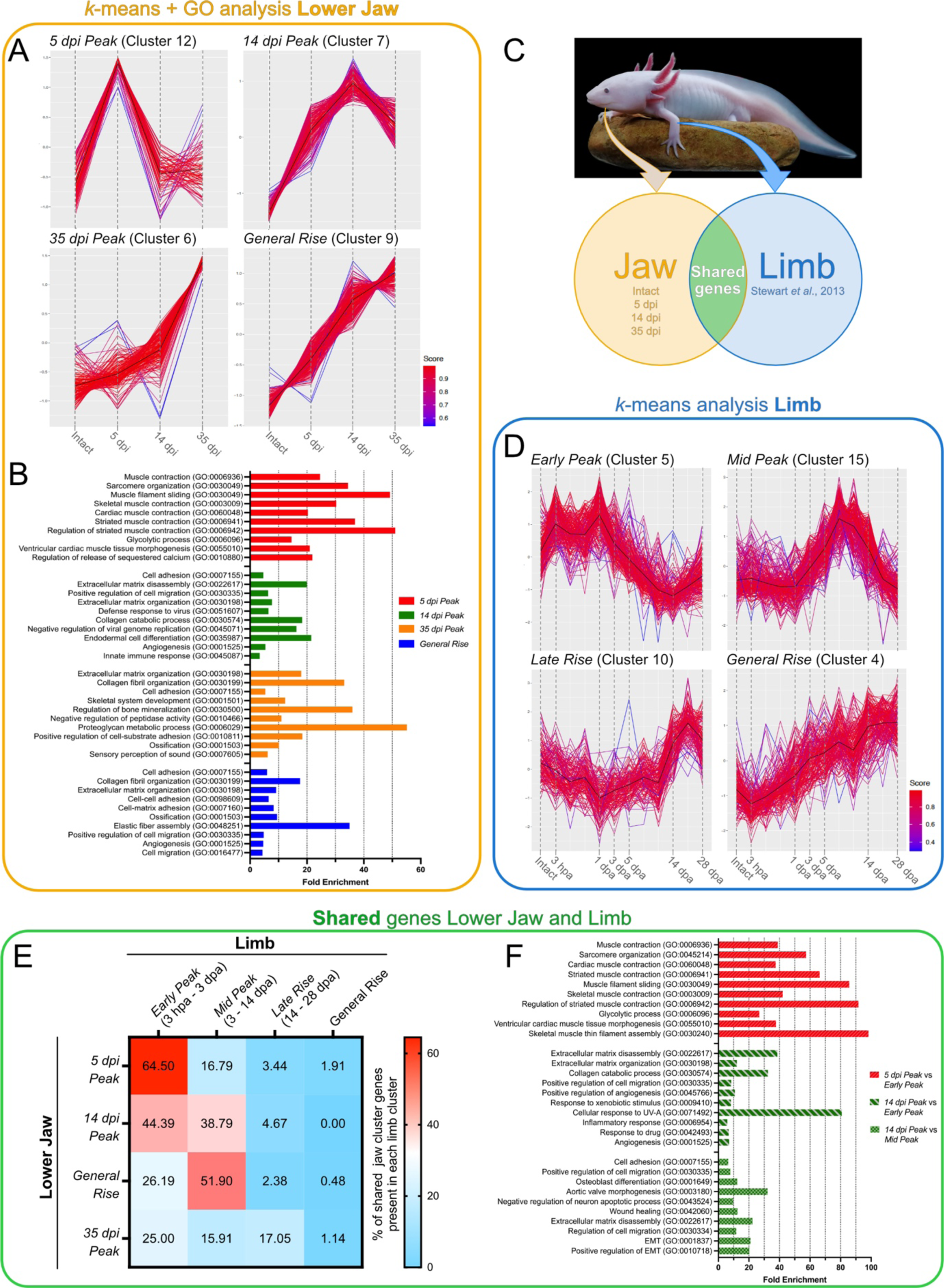
Regenerating mandibles after lateral resections exhibit transcriptomic similarities to limb regeneration, but progress at a slower pace. A. Four representative *k*-means clusters depicting *5 dpi Peak*, *14 dpi Peak*, *35 dpi Peak* and *General Rise* phases of mandible regeneration after lateral resection. **B.** Top 10 significantly enriched GO terms of *5 dpi Peak*, *14 dpi Peak*, *35 dpi Peak* and *General Rise* phases. **C.** Schematic representation of time points and datasets used for the comparisons between mandible and limb regeneration. Axolotl shown is Aires & Keeley’s 3-year-old pet axolotl, *Normando*. **D.** Four representative *k*-means clusters depicting *Early Peak* (3 hpa-3 dpa), *Mid Peak* (3-14 dpa), *Late Rise* (14-28 dpa), and *General Rise* phases of limb regeneration. Data from Stewart et al., 2013. **E.** Heatmap exhibiting the percentage of mandible genes shared with the limb. Displayed values do not total 100%, as genes that do not fit into these specified categories are not shown. **F.** Top 10 significantly enriched GO terms of the specified shared genes. Enriched GO terms in B and F are sorted by increasing p-value.

We then subjected the genes in each of these 4 groupings to gene ontology (GO) analysis to gain a better insight of the biological processes occurring during these four phases of regeneration (Fig. 3B, Table S3). This revealed that genes belonging to the group *5 dpi Peak* were mostly enriched in terms generally related to muscle contraction (expressing genes like *TnnI1*, *TnnI2*, *MyH1*) and sarcomere organization (e.g. *Tpm1*, *AnkrD1*, *MyoZ1*). In contrast, terms enriched in *14 dpi Peak* were mostly related to processes such as cell adhesion (*Itgb2*, *Itga5*, *Thbs1*, *Thbs4*), ECM disassembly (*Mmp1*, *Mmp9*, *Mmp10*, *Mmp13*), cell migration (*Tgfb1*, *Wnt5b*, *Mdk*, *Snai1*) and ECM organization (*Kazald2*, *Adamts4*, *Col7a1*, *Matn2*). Remarkably, there were signs of an ongoing immune response even two weeks after the injury, as GO terms associated to defense against viruses (*Rtp4*, *Ifitm1*, *RnaseL*, *Mx1*) and the innate immune response (*Arg1*, *Mpeg1*, *Treml1*, *Marco*, *Csf1r*) were found to be enriched in *14 dpi Peak*. In the *35 dpi Peak* group, besides a continuing prevalence of terms related to ECM organization, we also saw terms specifically associated to collagen fibril organization (*Lum*, *Fmod*, *Col2a1*, *Col9a1*), skeletal system development (*Acan*, *Cnmd*, *Pth1r*, *Hapln1*), and regulation of bone mineralization and ossification (*Phospho1*, *Omd*, *Matn1*, *Col11a1*), which matched with the cartilage formation and intramembranous ossification detected at this time by histological analysis. Finally, most of the terms enriched in *General Rise* were similar to *35 dpi Peak*, being predominantly related to cell adhesion, collagen fibril organization, ECM organization and ossification. However, we noted that both angiogenesis (*Pdgfrb*, *Acvrl1*, *Sox18*, *Hspg2*) and elastic fiber assembly (*Fbl5*, *Ltbp3*, *Ltbp4*, *Efemp2*) were also enriched terms in our dataset, suggesting the progressive regeneration of blood vessels in the affected region.

We next asked how transcriptionally similar regeneration of the laterally resected mandible was to the regenerating limb, one of the most studied structures in the axolotl (Fig. 3C). To investigate that, RNA-Seq read data following long-term limb regeneration in axolotls was obtained from published data (Stewart et al., 2013). After remapping to the current axolotl genome, data from the limb was handled in the same manner as the jaw and was likewise subjected to a *k*-means clustering algorithm, which subset genes into 23 clusters based on their expression over time (Fig. S5, Table S1). Filtration of the genes following clustering left us with 5,163 genes of interest, more than double the amount identified in the jaw. Interestingly, although the number of genes of interest identified in both structures varied widely from each other, almost 70% (1,458 out of 2,134 genes) of the lower jaw were shared with the limb (Table S2.).

As in the lower jaw, subsets of these clusters grouped together into broader patterns of gene expression. For ease of comparison, 17 clusters were chosen that could be categorized into four groups corresponding to broad regeneration phases: *Early Peak*, comprising clusters with peaks in the early phases of regeneration (3 hpa-3 dpa); *Mid Peak*, whose clusters peak during the intermediate phases of limb regeneration (3-14 dpa); *Late Rise*, containing clusters that peak at the later stages of limb regeneration (14-28 dpa); and *General Rise*, whose clusters demonstrate a steady increase over time (Fig 3D, Fig. S6). We next examined the transcriptomic similarities between mandible and limb in these four regeneration phases by assessing the amount of genes within each lower jaw category that were present in each limb grouping, which we designated as shared genes (Fig. 3E, Table S2). This approach revealed that 64.50% of the total shared genes in the *5 dpi Peak* jaw group were found in the *Early Peak* limb group. This indicated that the expression of many genes within the first days of regeneration was common to both mandible and limb regeneration, and GO analysis found that these genes were mostly associated with muscle contraction (Fig. 3F, Table S4). Notably, 59.76% of the shared genes between jaw *5 dpi Peak* and limb *Early Peak* actually peaked at 3 hpa in the limb, and then returned to basal levels or lower by 1 dpa (Table S2). Thus, their presence at 5 dpi in the mandible indicates that many genes associated with the initial wound response are present for a longer time in the lower jaw.

Interestingly, the *14 dpi Peak* mandible group displayed high overlap with different limb groupings, as 44.39% and 38.79% of shared genes in this set were present in limb *Early Peak* and *Mid Peak*, respectively (Fig. 3E). GO analysis showed that, in both cases, there was an enrichment in genes related to processes of cell migration, cell adhesion, ECM disassembly and ECM organization (Fig. 3F, Table S4), all of which previously associated with the formation of a blastema (reviewed in Aires et al., 2020). Yet, there were noticeable differences in the composition of the shared jaw genes within each of the two limb groups. Specifically, the shared genes between jaw *14 dpi Peak* and limb *Early Peak* showcased a higher diversity of matrix metalloproteinases (*Mmp1*, *Mmp3*, *Mmp9*, *Mmp10*, *Mmp13*), compared to only two (*Mmp11* and *Mmp13*) existing in the limb *Mid Peak*. The former thus correlates more closely with the earlier stages of limb regeneration, in which histolysis mostly occurs (Vinarsky et al., 2005; Yang and Bryant, 1994). Additionally, this group of shared genes was also enriched in factors related to the immune and inflammatory responses, which are likewise typical of early limb regeneration. In contrast, the jaw *14 dpi Peak* genes shared with the limb *Mid Peak* were particularly enriched in terms linked to mid-to-later stages of limb regeneration, such as cell proliferation (*Tgfb1*, *Mdk*, *Thbs4*), epithelial-to-mesenchymal transition (*Snai1*, *Hgf*, *Flna*), and osteoblast differentiation (*Vcan*, *Cbfb*, *Lox*, *Spp1*, *Tnc*) (Fig. 3F, Table S4). Moreover, we saw that *Kazald2* (previously identified as *Kazald1* in Bryant et al., 2017, see Materials and Methods) was present among these shared genes, which is specifically expressed in the limb blastema. Therefore, these observations suggested two concurrent stages of regeneration taking place in the 14 dpi lower jaw: an earlier phase in which the immune response and histolysis are still occurring, and a later one characterized by the presence of a blastema, as well as cartilage and bone formation. Indeed, this was supported by our histological evidence at this time point, as the distal stump already exhibited robust cartilage growth whereas none was yet observed in the proximal stump. It may also suggest that regeneration in at least a part of the mandible is starting to progress at a slower pace than in the limb.

This idea was further supported by the surprising find that 51.90% of shared genes in the *General Rise* mandible group were present in the *Mid Peak* limb group, instead of the expected *Late Rise* or *General Rise* groupings. Additionally, the low percentage of shared genes at late stage jaw and limb time points (*35 dpi Peak* and *General Rise* jaw groups vs. *Late Rise* and *General Rise* limb groups), likely reflects the fact that by this time the limb is becoming terminally differentiated, and thus not expressing genes that would be found in the mandible. While the lower jaw might also be expressing genes associated with differentiated tissues, the high amount of genes present that are shared with earlier phases in the limb imply that parts of the jaw are still actively regenerating at this time, and thus mandible regeneration is overall delayed compared to the limb.

Altogether, our results indicate that limb and laterally resected lower jaw regeneration share significant transcriptional similarities, especially in the early stages. Specifically, we could distinguish all the major stages typical of the regeneration of a complex structure in our model: earlier phases of muscle contraction-associated gene expression, immune system response, and tissue histolysis; processes involved in blastema formation, such as cell migration, proliferation and adhesion; and later stages of tissue differentiation, like cartilage and bone formation. However, our comparisons with the limb revealed mandible regeneration to proceed at a slower pace, especially during the mid-to-late stages, and that this rate can be different even in different regions within the regenerating structure.

### Regeneration of lateral mandible defects is associated to the formation of a WE/AEC and a blastema

Finally, we investigated if factors previously demonstrated to be important for blastema and WE/AEC formation during limb regeneration are likewise present during lower jaw regeneration. One marker particularly associated with the limb blastema is *Kazald2* (Bryant et al., 2017), which our analyses of limb RNA-Seq found to belong to cluster 15, and thus be part of the *Mid Peak* limb expression grouping (Fig. 3D). In the jaw, this gene fell into cluster 7, which was part of the *14 dpi Peak* grouping (Fig. 3A). Furthermore, in situ hybridization (ISH) analysis revealed that this gene was detected at 14 dpi in a salt-and-pepper pattern specifically in the mesenchymal cells composing part of the blastema-like structure (Fig. 4B). This similarity in expression pattern and timing confirmed that *Kazald2* expression is restricted to blastemal stages in both the regenerating limb and mandible.

**Fig.4.**
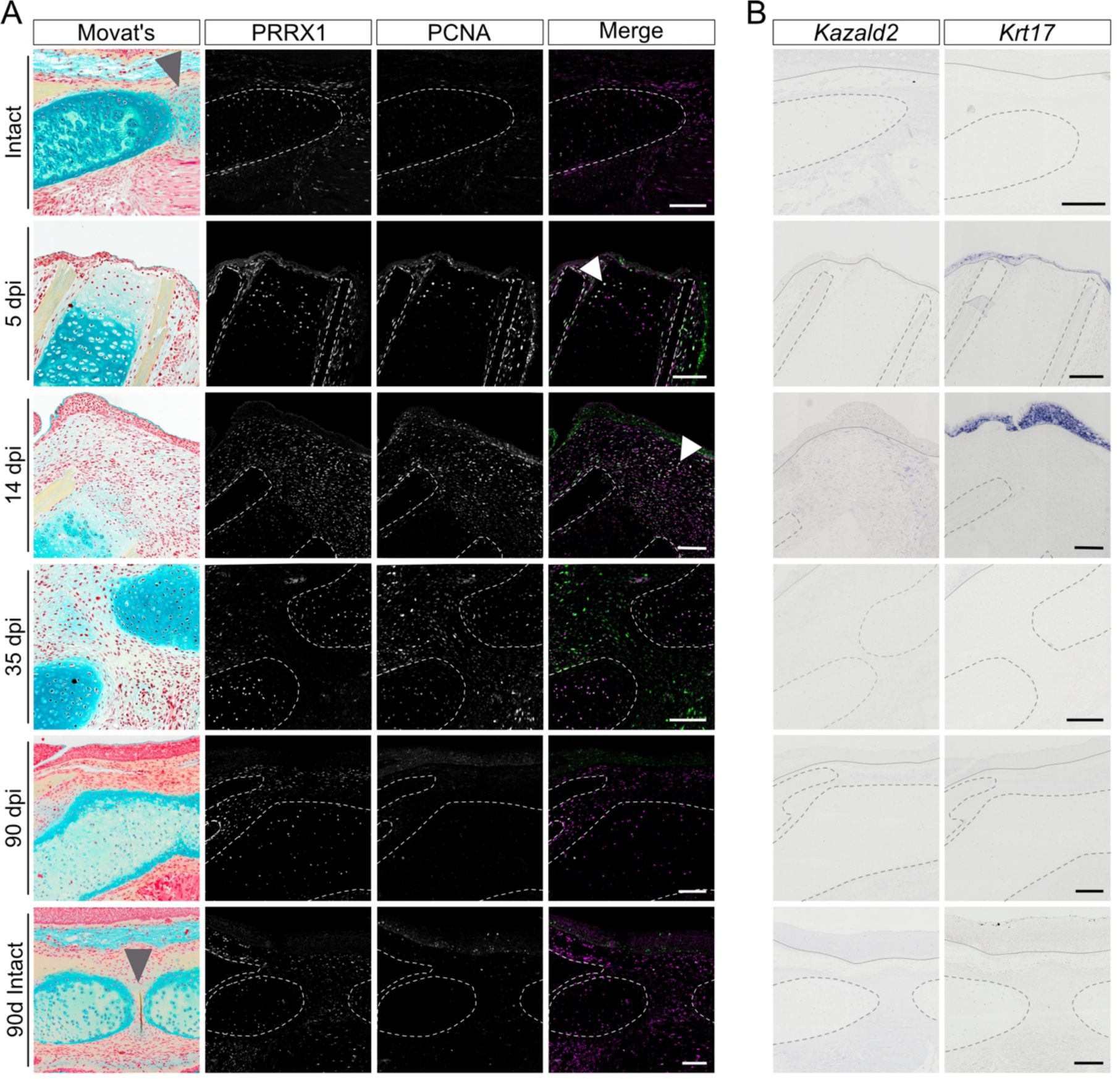
Regenerating mandibles express known markers involved in limb regeneration. A. Immunofluorescence (IF) of PRRX1 and PCNA in intact mandibles at the beginning (Intact) and end (90d Intact) of the experiment, and resected mandibles at 5-, 14-, 35-, and 90 dpi. All Movat’s pentachrome images are from insets in Fig.2 and Fig.S2, and are given here to provide tissue context. Gray arrowheads indicate the position of the medial symphysis in intact animals; white arrowheads show/highlights double positive cells. **B.** *In situ* hybridization (ISH) of *Kazald2* and *Krt17* in intact mandibles at the beginning (Intact) and end (90d Intact) of the experiment, and resected mandibles at 5-, 14-, 35-, and 90 dpi. Dotted lines show the outline of cartilage and ossified tissue, continuous lines indicate the border between WE/AEC and underlying mesenchyme. Scale bars: 250 µm.

Another gene of interest was *Krt17*, which was previously shown to be differentially expressed in the WE/AER during limb regeneration from early wound healing until palette stages (Leigh et al., 2018). Expression analysis by ISH in regenerating mandibles revealed that this gene was specifically detected in the epithelium covering the wound at 5 dpi, and was later found in the multi-layered epithelium at 14 dpi (Fig. 4B). This indicates that these two structures observed at 5- and 14 dpi most likely correspond to the limb’s WE and AEC, respectively. Furthermore, similar to limb regeneration in which *Krt17* was included in the *Mid Peak* group (cluster 2, Fig. S5, Fig. S6), *Krt17* was included in the *14 dpi Peak* jaw group (cluster 7, Fig. 3A), further emphasizing the similarities between regeneration of the two structures.

Finally, we examined *Prrx1* and *Pcna*. *Prrx1* is a transcription factor that is activated in dermal fibroblasts, which are major contributors to limb blastema formation (Satoh et al., 2007), while *Pcna* is a commonly utilized marker of cell proliferation (Miyachi et al., 1978). Immunofluorescence (IF) for PCNA revealed that intact lower jaws exhibited very little overall proliferation, and that PRRX1 was present in a variety of tissues, including the mandibular cartilage, mesenchyme, and cells of the medial symphysis (Fig. 4A, top and bottom rows). Following resection, by 5 dpi we observed an increase in proliferation, specifically in the epithelium adjacent to the injury, in the perichondrium, and in cells of the Meckel’s cartilage proximal to the resection plane. A proportion of these cells were also positive for PRRX1, particularly in the cartilage region that exhibited erosion of its ECM (Fig. 4A, white arrowhead). At 14 dpi, cell proliferation and PRRX1 presence dramatically increased, especially in the mesenchymal blastema-like structure, where we could detect many double positive cells (Fig. 4A, white arrowhead). We also saw considerable proliferation in the wound epithelium, which had developed an AEC-like appearance. By 35 dpi, both factors were still widely detected, but their co-expression substantially decreased. Proliferation was mostly observed in the connective tissue and nascent cartilage, whereas PRRX1 was chiefly present in more differentiated cartilage. Finally, at 90 dpi, PCNA and PRRX1 had mostly returned to intact levels. Importantly, the observed PRRX1 dynamics throughout jaw regeneration was consistent with our RNA-Seq data, which placed this gene in Cluster 9 of mandible regeneration, and thus part of its *General Rise* grouping (Fig. 3A). Meanwhile, in the limb, PRRX1 fell into cluster 15, which was part of its *Mid Peak* grouping (Fig. 3D). This further supports the view that the lower jaw regenerates at a slower pace than the limb, especially at later stages of regeneration.

In conclusion, morphological, histological, and transcriptomic data all suggest that, similar to the limb, laterally resected mandibles regenerate through the formation of a blastema and a WE/AEC that disappear in later stages of tissue differentiation. Moreover, expression analysis of important marker genes for the blastema and WE/AEC reinforces and validates our previously found transcriptomic parallels between mandible and limb regeneration in the axolotl. Specifically, the robust cell proliferation, and the presence of markers such as *Kazald2, Krt17,* and PRRX1 point towards a common underlying blastemal molecular signature in mandible and limb, despite them being formed in different regions in the body with distinct embryonic origins.

## Discussion

Symphyseal and parasymphyseal fractures of the mandible account for about one-third of all mandibular fractures (Atilgan et al., 2010; Rashid et al., 2013), since this region is particularly susceptible to trauma from falls and interpersonal violence, which is exacerbated by specific structural features that create predisposed breakage points. Apart from physical injuries, diseases such as cancer may frequently require the partial removal of the mandible and adjacent soft tissues in order to ensure the complete removal of the tumor (Eichberger et al., 2022). Thus, understanding how jaw repair could be improved has been the interest of much previous work.

Fundamental science has contributed to this field with several seminal studies in salamanders focusing on jaw regeneration. These mostly described its progression and positional limits (Ghosh et al., 1994; Goss and Stagg, 1958), the effects of fracture and cartilage formation (Hall and Hanken, 1985), and how specific features like the regeneration of the dental lamina occur(Graver, 1973; Graver, 1974; Graver, 1978). In this study, we add to this body of work by analyzing the regeneration of full-thickness lateral defects in the axolotl’s symphyseal and parasymphyseal regions, which are similar to defects in humans, and comparing its molecular profile to that of limb regeneration, as both structures are similar in their tissue complexity. This allowed us to discover morphological, histological and transcriptomic differences and similarities throughout the regeneration of these two structures.

### Two-phased gap minimization is a major event in lateral mandibular regeneration in the axolotl

Macroscopic analysis revealed that one of the major events during axolotl lower jaw regeneration following lateral resection was a two-phased minimization of the defect: the first phase involved rapid tissue contraction within the first day post-injury, whereas the second phase consisted of the displacement of the medial stump towards the defect during the second week of regeneration. Interestingly, displacement of the mandible stumps resulting in shifts of the midline was observed in previous studies of newt mandible regeneration, being reported either as a result of different rates of cartilage formation in lateral resections (Graver, 1978), or as a consequence of muscle retraction following distal transverse amputations (Ghosh et al., 1994; Goss and Stagg, 1958). A similar situation seems to also occur in humans following the fracture or loss of complete segments of the mandible, in which the direction of stump displacement depends on the localization of the defect in relation to the masticatory muscles. This can lead to either a convergence of the stumps and, thus, to defect reduction, or a divergence of the stumps and, consequently, to the enlargement of the defect region (Onodera et al., 2023). In our study, the observed gap minimization may have likewise resulted from the sudden mobility of the stumps due to severance of muscles or, alternatively, from a more active process directing these mechanical tissue readjustments. In fact, by 5 dpi, we observed an enrichment of GO terms associated with muscle contraction in the *5 dpi Peak* jaw group. This would not only contribute to the repair of the mandibular muscles, but could also drive the observed second phase of gap minimization in the mandible from 7 dpi onwards. Upregulation of muscle contraction-related genes was also described in early stages of resected newt and zebrafish mandible regeneration (Kurosaka et al., 2008; Ohgo et al., 2019), indicating that this could be a general feature of lower jaw regeneration. Given the mandible’s essential role in feeding behaviors (Lauder and Shaffer, 1985) minimization of the defect through muscle contraction would function as a “quick-fix” solution for the animal by providing a semi-functional lower jaw to catch prey as it undergoes regeneration. However, the importance of gap minimization processes, as well as their contribution to the speed of mandible regeneration, remains unknown.

### The axolotl mandible regenerates large defects by tissue growth and robust cartilage integration

The regenerative capabilities of the axolotl mandible after lateral resections are even more remarkable given the large size of the defect, which encompassed approximately 40% of one mandibular side. In contrast, segmental defects of this size in the axolotl limb skeletal elements fail to regenerate, and the injury develops a fibrotic response, in which the gap is instead filled with collagen fibers, connective tissue and regenerating musculature (Chen et al., 2015). Interestingly, even when these defects are small enough to elicit a regenerative response, the ends of the fragments are frequently misaligned, failing to integrate and heal adequately without external intervention (Chen et al., 2015; Polikarpova et al., 2022). This is similar to the human case, in which extensive jaw defects also show tight connective tissue scarring in the area of the missing mandibular segment (Lilly et al., 2021). One likely explanation for this proficiency to regenerate large defects in the lower jaw is the combination of gap minimization processes with formation of cartilage from both the proximal and distal mandibular stumps. Indeed, bilateral cartilage growth was also described in earlier work focusing on lateral resections in the newt mandible (Graver, 1978). Interestingly, this shows that, both in axolotl and newt, cartilage formation after lateral resections does not follow the rule of distal transformation (i.e., regenerating identities only distal to the amputation plane) as it does in the limb (Kragl et al., 2009). We have also noticed that, in axolotl, the regenerated cartilage was continuous with the previously existing Meckel’s cartilage in the proximal and distal stumps. This contrasted with adult newts, in which early chondrification in the proximal stump was generally initiated from and in direct contact with the prearticular bone (Ghosh et al., 1994; Goss and Stagg, 1958; Graver, 1978). Indeed, we detected proliferating chondrocytes close to the amputation plane at 5 dpi. This mode of cartilage regeneration would be another departure from the limb, in which new cartilage arises from the dedifferentiation and redifferentiation of fibroblasts into chondrocytes (Currie et al., 2016; Kragl et al., 2009).

Another important difference between limb and lower jaw regeneration is the process of bone formation, which follows their respective developmental modes of ossification (Olsen et al., 2000; Roberts and Hartsfield, 2004). In the mandible, we observed that the dentary bone was re-forming by intramembranous ossification from the proximal stump at 35 dpi, covering a significant portion of the new cartilage at 90 dpi. In contrast, post-regenerative ossification in the limb occurs through a cartilage intermediate, i.e. endochondral ossification (Olsen et al., 2000). Interestingly, we also found that the both coronoid and pre-articular bones had regenerated, as observed by the presence of new coronoidal teeth and the extension of new prearticular bone past the amputation plane. While the dentary bone is known to fully regenerate in both larval and adult newts, the prearticular bone, however, is able to regrow only in the former (Goss and Stagg, 1958; Graver, 1978). To our knowledge, this is the first example of prearticular bone regeneration in juvenile salamanders. Still, more work is needed to explore not only the origin of blastema cells, but also the modes of cartilage and bone regeneration in the mandible.

Finally, in our experiments, no medial symphysis was reformed even 90 days after resection. Absence of medial symphysis regeneration has also been sometimes reported after transverse amputations in several species of salamanders (Graver, 1973; Graver, 1974; Kurosaka et al., 2008), but other studies in newt have suggested that it can be entirely reconstituted (Ghosh et al., 1994; Goss and Stagg, 1958). The medial symphysis connects the two hemi-mandibles by fibrocartilage, which allows mobility between the bones while still connecting them. This structure is an important growth center of the mandible, persisting in most vertebrates except in mammals, in which it ossifies and becomes nearly undetectable (Biosse Duplan et al., 2016; Roberts and Hartsfield, 2004). In mammals, including humans, diseases or trauma injuring facial growth centers, can lead to severe growth retardation and deformities of the entire hemimandible (Cheong and Lo, 2011; Hofmann et al., 2023; Luo et al., 2016). Hence, the presence of the symphyseal region of the contralateral hemimandible in our resection models might have allowed cartilage formation to occur from its distal stump, but resulted in the absence of the medial symphysis. However, the role of growth centers in axolotl lower jaw regeneration is still unknown.

### Mandible and limb share a common regenerative program

Using the axolotl to study mandible regeneration allowed us to compare this process to that of the limb. Limb regeneration is, by far, the most well studied process in the axolotl, and its regenerative phases, as well as the molecular players involved, are relatively well known. Limb regeneration, thus, stands as the reference for the regeneration of any complex structure in the axolotl. Hence, our transcriptomic analysis enabled not only the characterization of the molecular mechanisms involved in mandible regeneration for the first time, but also to explore its similarities with the regenerating limb.

In general, the transcriptomic profile of the regenerating mandible at 5-, 14-, and 35 dpi fundamentally aligned with our histological observations. It also uncovered molecular signatures associated with the main stages of regeneration in the lower jaw, such as immune response, tissue histolysis, blastema formation, and chondrogenesis and ossification. Importantly, comparison to the limb revealed that the transcriptomic profiles of both structures overlapped greatly, with approximately 70% of genes expressed in mandible regeneration also being found during limb regeneration. Of these, the highest overlap occurred specifically in the blastema forming stages, which strongly indicates that the laterally resected mandible and limb share a common “blastema forming” molecular program. Indeed, we were able to validate these results by assessing the presence of three important limb regeneration markers: *Kazald2*, *Prrx1*, and *Krt17*. We confirmed that *Kazald2* and *Prrx1* were specifically found in our histologically-identified blastema-like structure, matching what occurs in the limb blastema (Bryant et al., 2017; Satoh et al., 2007). Likewise, we detected *Krt17* in the wound epithelium and AEC-like stratified epithelium of the regenerating mandible, just as it is during limb regeneration (Leigh et al., 2018).

Such similarities are somewhat remarkable, as these structures differ not only in their location, but also in their embryonic origins. However, the great parallels in tissue composition of these two structures, namely a significant proportion of skeletal elements composed of cartilage and bone, might be a potential explanation. Molecularly, this is reflected by the presence of *Prrx1* in the mesoderm and connective tissue during the development and regeneration of both structures (Leussink et al., 1995; Satoh et al., 2007). Another interesting possibility is that most of the shared genes could be related to the process of dedifferentiation itself, as studies in the limb showed that mature connective tissue cells, which compose the majority of the blastema (Dunis and Namenwirth, 1977; Kragl et al., 2009; Muneoka et al., 1986), dedifferentiate to revert to an embryonic limb bud-like phenotype during regeneration (Gerber et al., 2018). Ultimately, distinguishing between these two hypotheses will require more in-depth studies. Indeed, only by characterizing blastemas from different body structures and contrasting and comparing them will we be able to infer both general principles driving regeneration, as well as any specific regenerative constraints due to location or tissue composition.

### Mandible regeneration progresses at a slower pace than the limb

RNA-Seq analysis also suggested that mandible regeneration occurred at a slower pace than in the limb. In fact, we found that while a considerable proportion of genes belonging to the jaw *5 dpi Peak* were shared with the limb *Early Peak* group, we surprisingly observed that a majority of them were mostly found to peak at 3 hpa in the limb and return to basal levels by 1 dpa. The extended presence of these genes in the mandible could be due to a variety of factors, such as the larger wound surface area, potential reinjury upon mouth movement, or just inherent differences between the tissues that make up the two structures. Regardless of the cause, this delay was found to persist throughout the remainder of mandible regeneration.

A clear example of this is in the lower jaw *14 dpi Peak*. While there was a significant overlap of its blastema-associated genes with the limb *Mid Peak*, which was confirmed by expression of *Kazald2* and *Krt17* in regenerating jaws at 14 dpi, we did detect differences in regenerative timing in different regions of the regenerating mandible at this time point. Mainly, our transcriptomic analysis detected that the *14 dpi Peak* lower jaw group still contained many genes from the *Early Peak* limb group. Histologically, this was particularly evidenced by the fact that cartilage formation was more advanced in the distal stump than in the proximal stump at 14 dpi. The reasons for this timing discrepancy are currently unknown. However, resections through the distal side transversed the medial symphysis, which contains a significant number of PRRX1+ connective tissue cells in intact conditions. In mammals, the medial mandibular symphysis acts as an important growth center for interstitial cartilage in the skull (Roberts and Hartsfield, 2004). This raises the possibility that, due to their normal involvement in lower jaw growth, cells in the medial symphysis might remain primed and thus be able to be recruited more quickly for regeneration-associated growth than cells in the proximal stump, leading to different speeds of regeneration across the mandible.

Finally, the delay was most clearly evident in the late stages of regeneration, with many genes within the *Mid Peak* limb group found to still be rising at 35 dpi in the lower jaw. This was corroborated by the extensive presence of PRRX1 in regenerating jaws at 35 dpi, especially in the cartilage, which indicated that this factor had not yet returned to intact levels. Moreover, a recent study in limb regeneration conducted in similarly sized axolotls showed that all limb structures are fully re-formed by 38 dpa (Wells et al., 2021), contrasting with the lower jaw at this time point, which is still undergoing cartilage and bone formation.

Overall, these examples across different phases of regeneration in the mandible paint a picture of the mandible regenerating at a slower pace than the limb, especially as regeneration progresses to later stages, although the exact reasons for this delay will require more work to identify. However, while the mandible may regenerate slower than the limb, the regenerative program used is clearly shared with the limb, which highlights the existence of conserved underlying mechanisms to regenerate structures and tissues in the axolotl.

### Mandible regeneration after lateral resection is mostly complete after 90 days

Ultimately, despite the slower pace, we observed that the axolotl had regenerated all structures of the lower jaw by 90 dpi and, importantly, its mandibular skeletal elements and teeth. This agrees with previous studies showing that axolotls can fully regenerate their teeth 42 days after dentectomy (Makanae et al., 2020), and with the progression of mandible regeneration in the adult newt, in which mature teeth appear 8-10 weeks after amputation in the newly regenerated dentary bone (Goss and Stagg, 1958; Graver, 1978). This, in combination with the regrowth of all mandibular skeletal elements, indicates that overall regeneration of the axolotl mandible is mostly complete by 90 dpi, although the resected mandibles were still 9% smaller in perimeter than their uninjured counterparts. However, the fact that regenerating limbs in similarly sized axolotls can take up to 130 days to reach the size of the contralateral limb (Wells et al., 2021) suggests that resected mandibles at 90 dpi could still be undergoing a regenerative growth phase and thus, given enough time, they could eventually achieve the size of an uninjured mandible.

In conclusion, by laterally resecting the symphyseal and parasymphyseal regions of the axolotl mandible, we demonstrate that axolotl lower jaw regeneration occurs successfully through 2 rounds of gap minimization, followed by blastema formation, bilateral cartilage growth, complete defect bridging by integration of the regenerated tissue into the pre-existing one, and robust bone formation. Moreover, by investigating the gene expression profile of regenerating lower jaws and performing comparative analysis against the limb, we found that they both exhibit similar phases of regeneration, but that the mandible progresses at a slower pace. Finally, the great similarities between mandible and limb regeneration specifically in blastema stages hint at the existence of a common blastemal program that transcends differences in tissue locations and embryonic origins. This kind of comparative studies of diverse regenerating structures in the same model organism are crucial to identify core mechanisms underlying the re-formation of complex structures, and will most certainly shed light on the minimum requirements for a successful regeneration in vertebrates, as well as hint at the constraints limiting regeneration in mammals and humans.

## Materials and Methods

### Animal Husbandry

Husbandry and experimental procedures were performed according to the Animal Ethics Committee of the State of Saxony, Germany. Animals used were selected by their size (snout to tail = ST; snout to vent = SV).

Axolotl husbandry was performed in the CRTD axolotl facility using methodology adapted from Khattak et al., 2014. and according to the European Directive 2010/63/EU, Annex III, Table 9.1. Axolotls were kept in 18-19°C water in a 12 h light/12 h dark cycle and a room temperature of 20-22°C. Animals were housed in individual tanks categorized by a water surface (WS) area and a minimum water height (MWH). Axolotls of a size up to 5 cm SV were maintained in tanks with a WS of 180 cm^2^ and MWH of 4.5 cm. Axolotls up to 9 cm SV were maintained in tanks with a WS of 448 cm^2^ and MWH of 8 cm.

### Axolotl surgery

Experiments were performed on 16 12 cm ST long juvenile *d/d* axolotls, of which 10 animals were used for tissue sectioning and 6 animals were used for gene expression analysis. Axolotls were placed in agar dishes filled with 0.01% benzocaine solution. Once anesthetized, 5 mm of the mandible leftwards from the midline was removed. Following surgery, animals were checked every day. In this work, we used the nomenclature common in axolotl, distal for the proximity to the mandibular midline and proximal for the proximity to the mandibular joint. This differs from humans, where the adjacent region to the midline in the lower jaw is called mesial, whereas the area closer to the joint is called distal.

### Tissue collection

Tissue collection was performed by euthanizing animals by immersion in a lethal dosage of anesthesia (0.1% benzocaine). For paraffin embedding and sectioning, whole lower jaws were collected and fixed in 1x MEMFa (MOPS 0.1M pH 7.4, EGTA 2mM, MgSO4 × 7 H2O 1mM, 3.7% formaldehyde) for at least 1 week at 4°C. For RNA-Seq experiments, resected fragments after the initial surgery were used as intact condition. These, and the area of regenerated jaw corresponding to the initial surgery region at 5-, 14-, and 35dpi, were immediately flash frozen in liquid nitrogen and stored at −80°C until processed for RNA extraction.

### RNA extraction, Library Preparation and Sequencing

RNA extraction was performed using RNAeasy Mini Kit according to the manufacturer’s instructions. Samples were disrupted and homogenized using the NG010 Tissue Grinder Mixy Professional (NIPPON Genetics) in 600 µl of RLT Buffer containing β-mercaptoethanol (M625, Sigma). Extracted RNA was stored at −80 until processed for sequencing.

RNA sequencing libraries were prepared using NEBNext Ultra II Directional RNA Lib Prep (Biomek i7) with estimated fragment sizes of 300 - 400 bp. Poly-dT pull down enrichment of mRNA was performed before sequencing 101 bp paired-end reads on an Illumina NovaSeq 6000 (Illumina), generating between 37 and 57 million read pairs per sample. RNA-Seq raw data (fastq) has been deposited in NCBI under the Gene Expression Omnibus (GEO) accession code GSEXXXXX. Read data for axolotl limb regeneration over the course of 28 days was downloaded from www.axolomics.org (Axolotl Timecourse Filtered Reads) (Stewart et al., 2013).

### Read Mapping and Transcript Expression Analysis

Both our generated lower jaw reads and the downloaded limb fastq files were trimmed of adapter sequences and low quality bases via the programs cutadapt (https://journal.embnet.org/index.php/embnetjournal/article/view/200) and fastq_quality_filter from the FASTX-Toolkit (Hannon, G.J. (2010) FASTX-Toolkit. http://hannonlab.cshl.edu/fastx_toolkit), respectively. The reads were then mapped against the current axolotl reference genome (www.axolotl-omics.org, AmexG_v6.0-DD), using HISAT2 ver. 2.2.1(Pertea et al., 2015) with standard parameters and a known-splicesite-infile created from the current axolotl annotation file (AmexT_v47-AmexG_v6.0-DD.gtf) via the hisat2_extract_splice_sites.py command. Reads aligned with an average mapping rate of 96.02% for the jaw and 78.06% for the limb. Transcript quantification was then conducted using StringTie (Pertea et al., 2015) with standard parameters and the option of assembling novel transcripts. Finally, normalized CPM values for each sample were calculated using the Bioconductor package edgeR (Robinson et al., 2009) for R. Normalized gene counts (CPM) are provided in the Table S1.

### K-Means Analysis and Gene Ontology Analysis

Gene expression data over time in the limb was clustered into distinct groups through the use of a *k*-means clustering algorithm (Hartigan and Wong, 1979) available in R. To improve the generation of clusters, genes with a very low mean expression value (counts per million [CPM] < 0.8) and very low variation (sample variance < 0.4) were filtered out. Subsequently, due to the low number of replicates per tissue/sample/condition, a strict filtering process was applied. First, genes which never achieved a maximum expression value greater than 20 CPM were removed. Subsequently, a scaling multiplier for each gene based on their minimum expression value was calculated, and the genes with a ratio between their maximum and minimum CPM values that did not surpass it were removed. Following this filtering, remaining gene CPM values over time were converted into z-scores to standardize their changes in expression (Table S1). Optimal number of clusters were determined to be 23 for the limb and 15 for the jaw via calculation and assessment of the gap statistic (Tibshirani et al., 2001). Finally, genes that possessed a good cluster score (Pearson correlation coefficient > 0.75) were selected as our genes of interest. Lists of the upregulated and downregulated genes of interest were analyzed for significantly enriched Gene Ontology (GO) Terms via DAVID v6.8 (Huang et al., 2009) and are available in Tables S3 and S4.

### Time course imaging and defect area and angle measurements

Imaging was performed using a Zeiss Discovery.V20 stereomicroscope (Plan S 0.63). The images were imported into Image J 1.53t (http://imagej.nih.gov/ij) using the Bio-Formats plug-in 6.11.1 (https://docs.openmicroscopy.org/bio-formats/6.11.1/useres/imagej/features.html). For the relative defect size measurements, the length of the defect, full jaw perimeter, and hemimandible perimeter were measured using the segmental line tool, and the relative size was calculated. To measure the area of the defect over time, the polygon selection tool was used. To measure the observed fragment displacement, the angle tool was used to calculate the angle between the proximal endpoints of the mandible and the medial resection margin, according to the schematics in Fig. 1C. All measurements were repeated three times per animal and the average of the measured values was calculated.

For the quantification of defect area and angle of dislocation, 6 animals were used at all time points, except at 90 dpi where 3 were used. The area of the defect was considered zero once tissue continuity at the mandible edge was observed. For the quantification of full mandible perimeter, 3 resected animals and one intact animal were used.

Statistical analysis was performed using Prism9 (GraphPad Software, LLC) for macOS. To assess differences in area of defect a one-way ANOVA using Dunnet correction for multiple comparisons was performed between time points. To assess differences in distal stump displacement, a one-way ANOVA using a posthoc Tukey’s multiple comparisons test was used to assess statistical significance. P-values <0.05 were considered statistically significant.

### Sectioning and Histological stainings

Axolotl lower jaws were fixed in MEMFa and decalcified in 0.5 M EDTA for 3 weeks with daily changes of the solution. Sample embedding, sectioning and staining was performed by the CMCB Histology Facility, Dresden. Briefly, samples were dehydrated in a series of EtOH in RNase-free water until 100% EtOH, and then embedded in paraffin. Longitudinal sections of 4.5 µm were generated using a microtome. Movat’s Pentachrome (Morphisto Art-Nr:12057) staining was performed according to the manufacturer’s instructions. Imaging was performed using an Olympus OVK automated slide scanner system (UPLFLN 4x/0.13 or UPLSAPO 10x/0.40).

### In situ hybridization in tissue sections

*In situ* hybridization was used to assess gene expression in sectioned tissues. Probes for the axolotl *Kazald2* and *Krt17* were generated as previously described (Riquelme-Guzmán et al., 2022). Primer sequences for probe amplification were taken from Leigh et al., 2018(*Krt17*) and Bryant et al., 2017 (*Kazald1*). We reannotated as *Kazald2* the gene previously identified as *Kazald1* by using phylogenetic analysis conducted with aLeaves (Kuraku et al., 2013), which placed it within the clade containing actinopterygian (including zebrafish) *Kazald2*, instead of the clade containing mammalian *Kazald1*.

Briefly, slides were dewaxed in Roti-Histol (Carl Roth 6640) and rehydrated through a series of EtOH in RNase-free water. After one wash in PBS, slides were fixed in 4% PFA + 0.2% glutaraldehyde for 20 min, washed twice in PBS, and treated with proteinase K (10 μg/ml in PBS) at 37°C for 10 min. Slides were washed again in PBS, post-fixed in PFA 4%, and incubated with 0.1 N HCL for 15 min. After being rinsed in 0.1 M Triethanolamine (TEA) at pH 7.5, slides were incubated in a freshly prepared solution of TEA 0.1 M and 0.1 M Acetic Anhydride for 10 min. Next, slides were rinsed in PBS, then RNase-free water, and incubated in prewarmed Hybridization Solution (50% formamide, 5x SSC [3 M NaCl, 300 mM sodium citrate] (pH 5.5), 0.1% Tween 20, 50 μg/ml yeast tRNA, 100 μg/ml heparin, 1x Denhart’s, 0.1% CHAPS, 5 mM EDTA) for 1 h. Slides were next incubated with Hybridization Solution containing the RNA probe overnight at 65°C and then washed at 65°C the following day twice with prewarmed 5x SSC, 2x SSC, and 0.2x SSC for 30 min each wash. Tissue was then washed at room temperature (RT) with TNE buffer (10 mM Tris-HCl (pH 7.5), 500 mM NaCl, 1 mM EDTA), treated with RNase (20 μg/ml in TNE buffer) for 10 min. and washed again with TNE buffer. The tissue was equilibrated with MABT (100 mM Maleic acid, 150 mM NaCl, 0.1 % Tween 20), blocked with MABT/Block (MABT containing 1% blocking reagent (Roche #11096176001)) for 1 h at RT, and incubated with a 1:5000 dilution of alkaline phosphatase-conjugated anti-digoxigenin antibody (Roche 11093274910) in MABT/Block overnight at 4°C. After extensive washes with MABT at RT, slides were equilibrated in NTMT (100 mM Tris-HCL (pH 9.5), 50 mM MgCl2, 100 mM NaCl, 0.1% Tween-20), and developed at RT in BM Purple (Roche 11442074001). Reactions were stopped with PBS and fixed with 4% PFA overnight. Finally, slides were dehydrated in a series of EtOH in water, incubated 10 min in Roti-Histol, and mounted in Entellan for imaging using an Olympus OVK automated slide scanner system (UPLSAPO 10x/0.40).

### Immunofluorescence in tissue sections

Sections were first deparaffinized by incubating the slides for 10 min in Roti-Histol, and rehydrated through a series of EtOH in RNase-free water. After 3 washes in PBS 1X, sections were subjected to antigen retrieval for 30 min at 95°C in Sodium Citrate Buffer (10mM Sodium Citrate, 0.005% Tween-20, pH 6.0). Then, slides were allowed to cool down for 30 min before being washed 3 times with PBS 1X and blocked with blocking solution (PBS 1X + 0.3% Triton-X + 10% goat serum) for 1 hour. Primary antibody incubation of anti-PRRX1 (rabbit, MPI-CBG antibody facility and a kind gift from Prayag Murawala) and anti-PCNA (mouse) conjugated with Alexa 647 (Santa Cruz PC10, sc-56 AF647) was done in a blocking solution 1:100 overnight at 4°C. The following day, sections were washed 3 times with PBS 1X for 10 min and incubated with secondary antibody goat anti-rabbit Alexa Fluor 568 (Invitrogen A-11011) in blocking solution for 3 hours at RT. Finally, sections were washed again 3 times in PBS 1X, incubated with Hoechst 33258 (Abcam Ab228550) 1:1000 in PBS 1X for 10 min, rinsed in PBS 1X and mounted in VectaShield® (Vector Laboratories, H-1000-10). Imaging was performed using a Zeiss confocal laser scanning microscope LSM 780 (Plan-apochromat 10x/0.45).

### Image processing and analysis

All images were processed using Fiji (Schindelin et al., 2012). Processing involved selecting regions of interest, merging, or splitting channels and improving brightness levels for proper presentation in figures. IF and ISH were performed on sister slides through the same region of the same sample.

## Acknowledgments

We would like to thank past and current members of the Sandoval-Guzmán lab for continuous support and companionship during the development of this work. We are also grateful to Anja Wagner, Beate Gruhl, and Judith Konantz for their dedication to the axolotl care. This work was supported by the Light Microscopy Facility, the Histology Facility, and the Genome Facility, all of them Core Facilities of the CMCB Technology Platform at TU Dresden. R.A was supported by a Von Humboldt Foundation Research fellowship PRT 1208176 HFST-P and a DFG Eigene Stelle Grant AI 214/1-1. SDK was supported by the DIGS-BB graduate program, Dresden.

## Author contributions

Conceptualization: mainly R.A., J.K.,

Methodology: R.A., J.K., S.D.K.

Investigation: R.A., J.K, S.D.K.

Project administration: T.S.G.

Supervision: T.S.G., G.L.

Funding acquisition: T.S.G., G.L., T.A.S

Data curation: R.A., S.D.K.

Writing – original draft: mainly R.A., J.K.,

Writing – review and editing: all authors.

## References

1. Aires, R., Keeley, S. D. and Sandoval-Guzmán, T. (2020). Basics of Self-Regeneration. In Cell Engineering and Regeneration (ed. Gimble, J. M.), Marolt Presen, D.), Oreffo, R.), Redl, H.), and Wolbank, S.), p. Cham: Springer International Publishing.

2. Atilgan, S., Erol, B., Yaman, F., Yilmaz, N. and Ucan, M. C. (2010). Mandibular fractures: a comparative analysis between young and adult patients in the southeast region of Turkey. Journal of Applied Oral Science 18, 17–22.

3. Atkins, J. B., Houle, L., Cantelon, A. S. and Maddin, H. C. (2020). Normal development in Ambystoma mexicanum: A complementary staging table for the skull based on Alizarin red S staining. Developmental Dynamics 249, 656–665.

4. Biosse Duplan, M., Komla-Ebri, D., Heuzé, Y., Estibals, V., Gaudas, E., Kaci, N., Benoist-Lasselin, C., Zerah, M., Kramer, I., Kneissel, M., et al. (2016). Meckel’s and condylar cartilages anomalies in achondroplasia result in defective development and growth of the mandible. Hum Mol Genet ddw153.

5. Bräuer, C., Ullmann, K., Lauer, G., Franke, A., McLeod, N. M. H. and Leonhardt, H. (2023). Alloplastic reconstruction of the mandible after subtotal mandibulectomy for medication-related osteonecrosis of the jaw: An update of the method. Head Neck 45, 2638–2648.

6. Bryant, D. M., Johnson, K., DiTommaso, T., Tickle, T., Couger, M. B., Payzin-Dogru, D., Lee, T. J., Leigh, N. D., Kuo, T.-H., Davis, F. G., et al. (2017). A Tissue-Mapped Axolotl De Novo Transcriptome Enables Identification of Limb Regeneration Factors. Cell Rep 18, 762–776.

7. Charbonneau, A. M., Roy, S. and Tran, S. D. (2016). Oral-Facial Tissue Reconstruction in the Regenerative Axolotl. J Exp Zool B Mol Dev Evol 326, 489–502.

8. Charbonneau, A. M., Åström, P., Salo, T., Roy, S. and Tran, S. D. (2021). Axolotls’ and Mices’ Oral-Maxillofacial Trephining Wounds Heal Differently. Cells Tissues Organs 210, 260–274.

9. Chen, X., Song, F., Jhamb, D., Li, J., Bottino, M. C., Palakal, M. J. and Stocum, D. L. (2015). The axolotl fibula as a model for the induction of regeneration across large segment defects in long bones of the extremities. PLoS One 10, 1–26.

10. Cheong, Y.-W. and Lo, L.-J. (2011). Facial asymmetry: etiology, evaluation, and management. Chang Gung Med J 34, 341–51.

11. Cohen, M. and Schultz, R. C. (1985). Mandibular Reconstruction. Clin Plast Surg 12, 411–422.

12. Currie, J. D., Kawaguchi, A., Traspas, R. M., Schuez, M., Chara, O. and Tanaka, E. M. (2016). Live Imaging of Axolotl Digit Regeneration Reveals Spatiotemporal Choreography of Diverse Connective Tissue Progenitor Pools. Dev Cell 39, 411–423.

13. Davit-Béal, T., Chisaka, H., Delgado, S. and Sire, J. (2007). Amphibian teeth: current knowledge, unanswered questions, and some directions for future research. Biological Reviews 82, 49–81.

14. Disa, J. J. and Cordeiro, P. G. (2000). Mandible reconstruction with microvascular surgery. Semin Surg Oncol 19, 226–234.

15. Dunis, D. A. and Namenwirth, M. (1977). The role of grafted skin in the regeneration of X-irradiated axolotl limbs. Dev Biol 56, 97–109.

16. Echeverri, K., Fei, J. and Tanaka, E. M. (2022). The Axolotl’s journey to the modern molecular era. In Current Topics in Developmental Biology, pp. 631–658. Academic Press Inc.

17. Eichberger, J., Weber, F., Spanier, G., Gerken, M., Schreml, S., Schulz, D., Fiedler, M., Ludwig, N., Bauer, R. J., Reichert, T. E., et al. (2022). Loss of MMP-27 Predicts Mandibular Bone Invasion in Oral Squamous Cell Carcinoma. Cancers (Basel) 14, 1–16.

18. Gerber, T., Murawala, P., Knapp, D., Masselink, W., Schuez, M., Hermann, S., Gac-Santel, M., Nowoshilow, S., Kageyama, J., Khattak, S., et al. (2018). Single-cell analysis uncovers convergence of cell identities during axolotl limb regeneration. Science (1979) 362,.

19. Ghosh, S., Thorogood, P. and Ferretti, P. (1994). Regenerative capability of upper and lower jaws in the newt. International Journal of Developmental Biology 38, 479–490.

20. Gilbert, S. F. (1997). *Developmental Biology*. Sunderland, Massachusetts: Sinauer Associates.

21. Goss, R. J. and Stagg, M. W. (1958). Regeneration of lower jaws in adult newts. J Morphol 102, 289– 309.

22. Graver, H. T. (1973). The polarity of the dental lamina in the regenerating salamander jaw. J Embryol Exp Morphol 30, 635–646.

23. Graver, H. T. (1974). Origin of the dental lamina in the regenerating salamander jaw. Journal of Experimental Zoology 189, 73–83.

24. Graver, H. T. (1978). Re-regeneration of lower jaws and the dental lamina in adult urodeles. J Morphol 157, 269–279.

25. Hall, B. K. and Hanken, J. (1985). Repair of fractured lower jaws in the spotted salamander: Do amphibians form secondary cartilage? Journal of Experimental Zoology 233, 359–368.

26. Hartigan, J. A. and Wong, M. A. (1979). Algorithm AS 136: A K-Means Clustering Algorithm. Appl Stat 28, 100.

27. Hofmann, E., Koerdt, S., Heiland, M., Raguse, J.-D. and Voss, J. O. (2023). Pediatric Maxillofacial Trauma: Insights into Diagnosis and Treatment of Mandibular Fractures in Pediatric Patients. Int J Clin Pediatr Dent 16, 499–509.

28. Huang, D. W., Sherman, B. T. and Lempicki, R. A. (2009). Systematic and integrative analysis of large gene lists using DAVID bioinformatics resources. Nat Protoc 4, 44–57.

29. Kakarala, K., Shnayder, Y., Tsue, T. T. and Girod, D. A. (2018). Mandibular reconstruction. Oral Oncol 77, 111–117.

30. Khattak, S., Murawala, P., Andreas, H., Kappert, V., Schuez, M., Sandoval-Guzmán, T., Crawford, K. and Tanaka, E. M. (2014). Optimized axolotl (Ambystoma mexicanum) husbandry, breeding, metamorphosis, transgenesis and tamoxifen-mediated recombination. Nat Protoc 9, 529–540.

31. Kragl, M., Knapp, D., Nacu, E., Khattak, S., Maden, M., Epperlein, H. H. and Tanaka, E. M. (2009). Cells keep a memory of their tissue origin during axolotl limb regeneration. Nature 460, 60–65.

32. Kuraku, S., Zmasek, C. M., Nishimura, O. and Katoh, K. (2013). aLeaves facilitates on-demand exploration of metazoan gene family trees on MAFFT sequence alignment server with enhanced interactivity. Nucleic Acids Res 41, W22–W28.

33. Kurosaka, H., Takano-Yamamoto, T., Yamashiro, T. and Agata, K. (2008). Comparison of molecular and cellular events during lower jaw regeneration of newt (Cynops pyrrhogaster) and West African clawed frog (Xenopus tropicalis). Developmental Dynamics 237, 354–365.

34. Lauder, G. V. and Shaffer, H. B. (1985). Functional morphology of the feeding mechanism in aquatic ambystomatid salamanders. J Morphol 185, 297–326.

35. Leigh, N. D., Dunlap, G. S., Johnson, K., Mariano, R., Oshiro, R., Wong, A. Y., Bryant, D. M., Miller, B. M., Ratner, A., Chen, A., et al. (2018). Transcriptomic landscape of the blastema niche in regenerating adult axolotl limbs at single-cell resolution. Nat Commun 9,.

36. Leussink, B., Brouwer, A., El Khattabi, M., Poelmann, R. E., Gittenberger-de Groot, A. C. and Meijlink, F. (1995). Expression patterns of the paired-related homeobox genes MHox Prx1 and S8 Prx2 suggest roles in development of the heart and the forebrain. Mech Dev 52, 51–64.

37. Lilly, G. L., Petrisor, D. and Wax, M. K. (2021). Mandibular rehabilitation: From the Andy Gump deformity to jaw-in-a-day. Laryngoscope Investig Otolaryngol 6, 708–720.

38. Luo, E., Du, W., Li, J., Zhu, S., Li, J. and Hu, J. (2016). Guideline for the Treatment of Condylar Osteochondroma Combined With Secondary Dentofacial Deformities. Journal of Craniofacial Surgery 27, 1156–1161.

39. Makanae, A., Tajika, Y., Nishimura, K., Saito, N., Tanaka, J. and Satoh, A. (2020). Neural regulation in tooth regeneration of Ambystoma mexicanum. Sci Rep 10, 9323.

40. Markwardt, J., Sembdner, P., Lesche, R., Jung, R., Spekl, K., Mai, R., Schulz, M. C. and Reitemeier, B. (2014). Experimental findings on customized mandibular implants in Göttingen minipigs - A pilot study. International Journal of Surgery 12, 60–66.

41. Miyachi, K., Fritzler, M. J. and Tan, E. M. (1978). Autoantibody to a nuclear antigen in proliferating cells. J Immunol 121, 2228–34.

42. Muneoka, K., Fox, W. F. and Bryant, S. V. (1986). Cellular contribution from dermis and cartilage to the regenerating limb blastema in axolotls. Dev Biol 116, 256–260.

43. Nauth, A., Schemitsch, E., Norris, B., Nollin, Z. and Watson, J. T. (2018). Critical-Size Bone Defects: Is There a Consensus for Diagnosis and Treatment? J Orthop Trauma 32, S7–S11.

44. Neufeld, D. A. and Aulthouse, A. L. (1986). Association of mesenchyme with attenuated basement membranes during morphogenetic stages of newt limb regeneration. American Journal of Anatomy 176, 411–421.

45. Ohgo, S., Ichinose, S., Yokota, H., Sato-Maeda, M., Shoji, W. and Wada, N. (2019). Tissue regeneration during lower jaw restoration in zebrafish shows some features of epimorphic regeneration. Dev Growth Differ 61, 419–430.

46. Olsen, B. R., Reginato, A. M. and Wang, W. (2000). Bone Development. Annu Rev Cell Dev Biol 16, 191–220.

47. Onodera, K., Miyamoto, I., Hoshi, I., Kawamata, S., Takahashi, N., Shimazaki, N., Kondo, H. and Yamada, H. (2023). Towards Optimum Mandibular Reconstruction for Dental Occlusal Rehabilitation: From Preoperative Virtual Surgery to Autogenous Particulate Cancellous Bone and Marrow Graft with Custom-Made Titanium Mesh—A Retrospective Study. J Clin Med 12, 1122.

48. Payzin-Dogru, D. and Whited, J. L. (2018). An integrative framework for salamander and mouse limb regeneration. International Journal of Developmental Biology 62, 393–402.

49. Pertea, M., Pertea, G. M., Antonescu, C. M., Chang, T. C., Mendell, J. T. and Salzberg, S. L. (2015). StringTie enables improved reconstruction of a transcriptome from RNA-seq reads. Nat Biotechnol 33, 290–295.

50. Polikarpova, A., Ellinghaus, A., Schmidt-Bleek, O., Grosser, L., Bucher, C. H., Duda, G. N., Tanaka, E. M. and Schmidt-Bleek, K. (2022). The specialist in regeneration—the Axolotl—a suitable model to study bone healing? NPJ Regen Med 7, 35.

51. Rashid, A., Eyeson, J., Haider, D., Van Gijn, D. and Fan, K. (2013). Incidence and patterns of mandibular fractures during a 5-year period in a London teaching hospital. British Journal of Oral and Maxillofacial Surgery 51, 794–798.

52. Reitemeier, B., Schöne, C., Lesche, R., Lauer, G., Schulz, M. C. and Markwardt, J. (2016). Contour identical implants to bridge mandibular continuity defects - individually generated by LaserCUSING® - A feasibility study in animal cadavers. Head Face Med 12, 1–8.

53. Riquelme-Guzmán, C., Tsai, S. L., Carreon Paz, K., Nguyen, C., Oriola, D., Schuez, M., Brugués, J., Currie, J. D. and Sandoval-Guzmán, T. (2022). Osteoclast-mediated resorption primes the skeleton for successful integration during axolotl limb regeneration. Elife 11, 2022.04.27.489662.

54. Roberts, W. E. and Hartsfield, J. K. (2004). Bone development and function: genetic and environmental mechanisms. Semin Orthod 10, 100–122.

55. Robinson, M. D., McCarthy, D. J. and Smyth, G. K. (2009). edgeR: A Bioconductor package for differential expression analysis of digital gene expression data. Bioinformatics 26, 139–140.

56. Satoh, A., Gardiner, D. M., Bryant, S. V. and Endo, T. (2007). Nerve-induced ectopic limb blastemas in the axolotl are equivalent to amputation-induced blastemas. Dev Biol 312, 231–244.

57. Schemitsch, E. H. (2017). Size Matters: Defining Critical in Bone Defect Size! *J Orthop Trauma* 31, S20–S22.

58. Schindelin, J., Arganda-Carreras, I., Frise, E., Kaynig, V., Longair, M., Pietzsch, T., Preibisch, S., Rueden, C., Saalfeld, S., Schmid, B., et al. (2012). Fiji: an open-source platform for biological-image analysis. Nat Methods 9, 676–682.

59. Simon, A. and Tanaka, E. M. (2013). Limb regeneration. Wiley Interdiscip Rev Dev Biol 2, 291–300.

60. Stewart, R., Rascón, C. A., Tian, S., Nie, J., Barry, C., Chu, L. F., Ardalani, H., Wagner, R. J., Probasco, M. D., Bolin, J. M., et al. (2013). Comparative RNA-seq Analysis in the Unsequenced Axolotl: The Oncogene Burst Highlights Early Gene Expression in the Blastema. PLoS Comput Biol 9,.

61. Tanaka, E. M. (2016). The Molecular and Cellular Choreography of Appendage Regeneration. Cell 165, 1598–1608.

62. Tibshirani, R., Walther, G. and Hastie, T. (2001). Estimating the number of clusters in a data set via the gap statistic. J R Stat Soc Series B Stat Methodol 63, 411–423.

63. Vinarsky, V., Atkinson, D. L., Stevenson, T. J., Keating, M. T. and Odelberg, S. J. (2005). Normal newt limb regeneration requires matrix metalloproteinase function. Dev Biol 279, 86–98.

64. Wells, K. M., Kelley, K., Baumel, M., Vieira, W. A. and McCusker, C. D. (2021). Neural control of growth and size in the axolotl limb regenerate. Elife 10,.

65. Yang, E. V. and Bryant, S. V. (1994). Developmental Regulation of a Matrix Metalloproteinase during Regeneration of Axolotl Appendages. Dev Biol 166, 696–703.

