## Supplemental Figures for "Axolotl mandible regeneration occurs through mechanical gap closure and a shared regenerative program with the limb"

### Contents:

**Supplementary Figure S1:** Complete time course of mandible regeneration in 3 individual animals

**Supplementary Figure S2:** Additional histological stainings at 5-, 14-, 35-, and 90 dpi.

**Supplementary Figure S3:** 15 clusters found by *k*-means clustering algorithm during lower jaw regeneration.

**Supplementary Figure S4:** Regenerating lower jaw cluster groupings for downstream analysis.

**Supplementary Figure S5:** 24 clusters found by *k*-means clustering algorithm during limb regeneration.

**Supplementary Figure S6:** Regenerating limb cluster groupings for downstream analysis.

**Table S1** Gene counts and clusters of mandible and limb.

**Table S2** Calculation of shared genes between mandible and limb at various time points.

**Table S3** Gene ontology analysis of *5 dpi Peak*, *14 dpi Peak*, *35 dpi Peak* and *General Rise* during lower jaw regeneration.

**Table S4** Gene ontology analysis of shared lower jaw regeneration cluster present in each limb regeneration cluster.

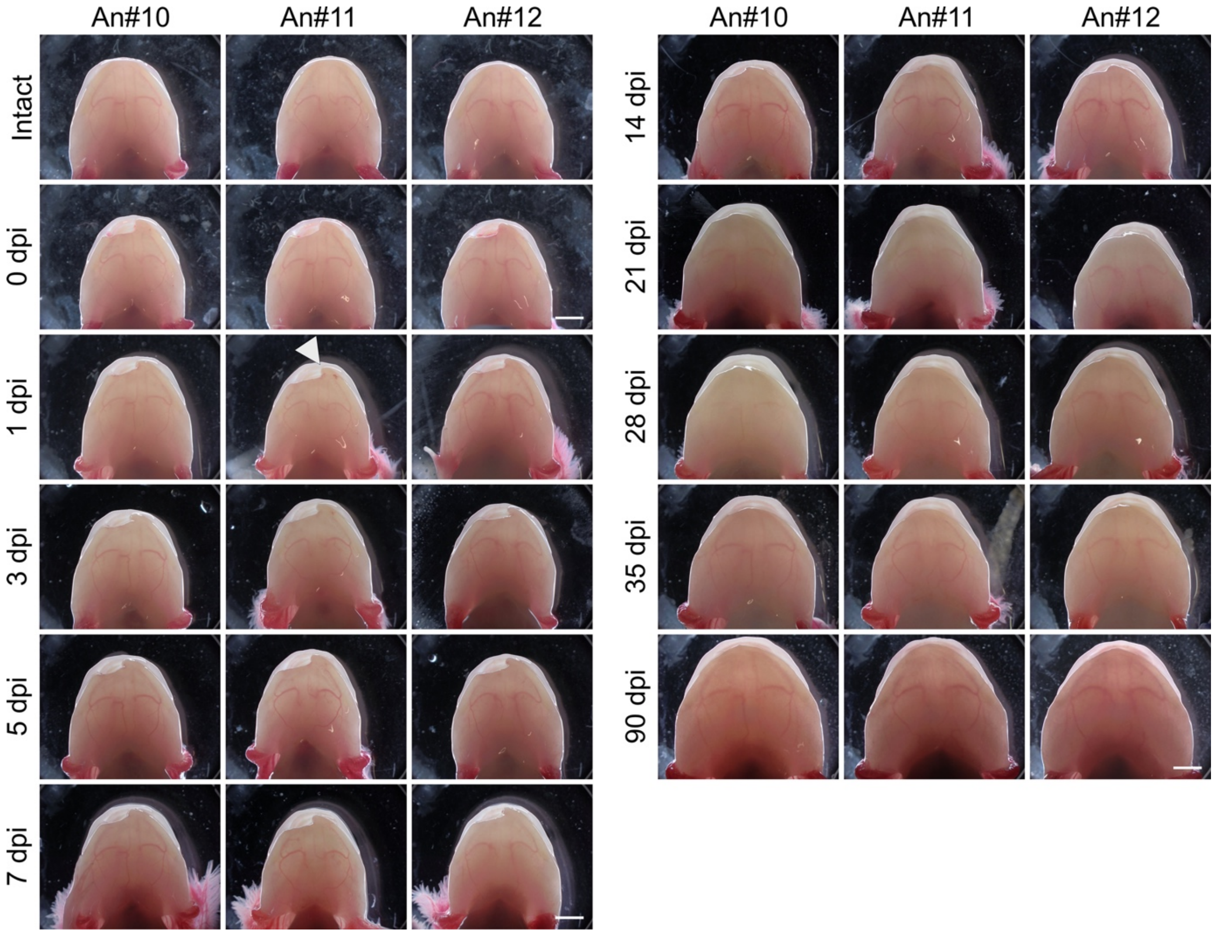


**Fig. S1: Time course of lower jaw regeneration after full-thickness lateral resections in 3 representative individuals**. Animals are pictured before (Intact) and immediately after resection (0 dpi), and at 1-, 3-, 5-, 7-, 14-, 21-, 28-, 35-, and 90 dpi. White arrowhead indicates medial edge of the defect. An#11 is used for time course in Fig.1. Scale bar: 5 mm.

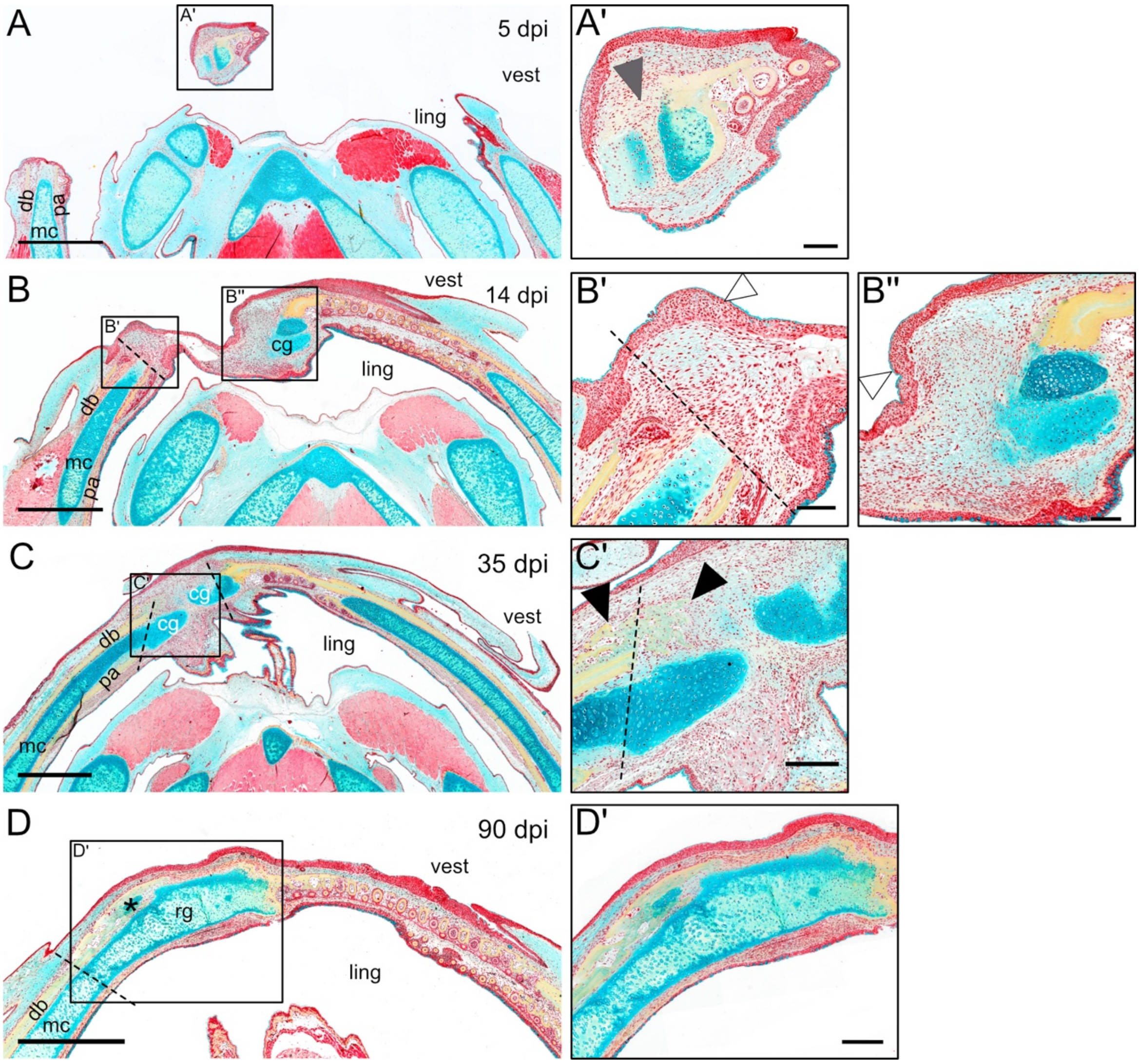


**Fig. S2: Movat's pentachrome staining of longitudinal sections of regenerating lower jaws after full-thickness lateral resections at 5- (A), 14- (B), 35- (C), 90 dpi (D).** Grey arrowhead in A' represents the medial symphysis. White arrowheads in B' and B" indicate the wound epithelium. Black arrowheads in C' show intramembranous ossification. Dashed lines indicate the approximate site of resection. mc: Meckel's cartilage, pa: prearticular, db: dentary; vest: vestibular side; ling: lingual side; cg: cartilaginous growth; rc: regenerated cartilage. Scale bar in A-D: 2 mm, scale bar in A', B', B", C': 250 µm. Scale bar in D': 500 µm. All of the sections correspond to different planes through the regenerated region in the same animals shown in Fig. 2. D corresponds to an intermediate section between Fig. 2E and F.

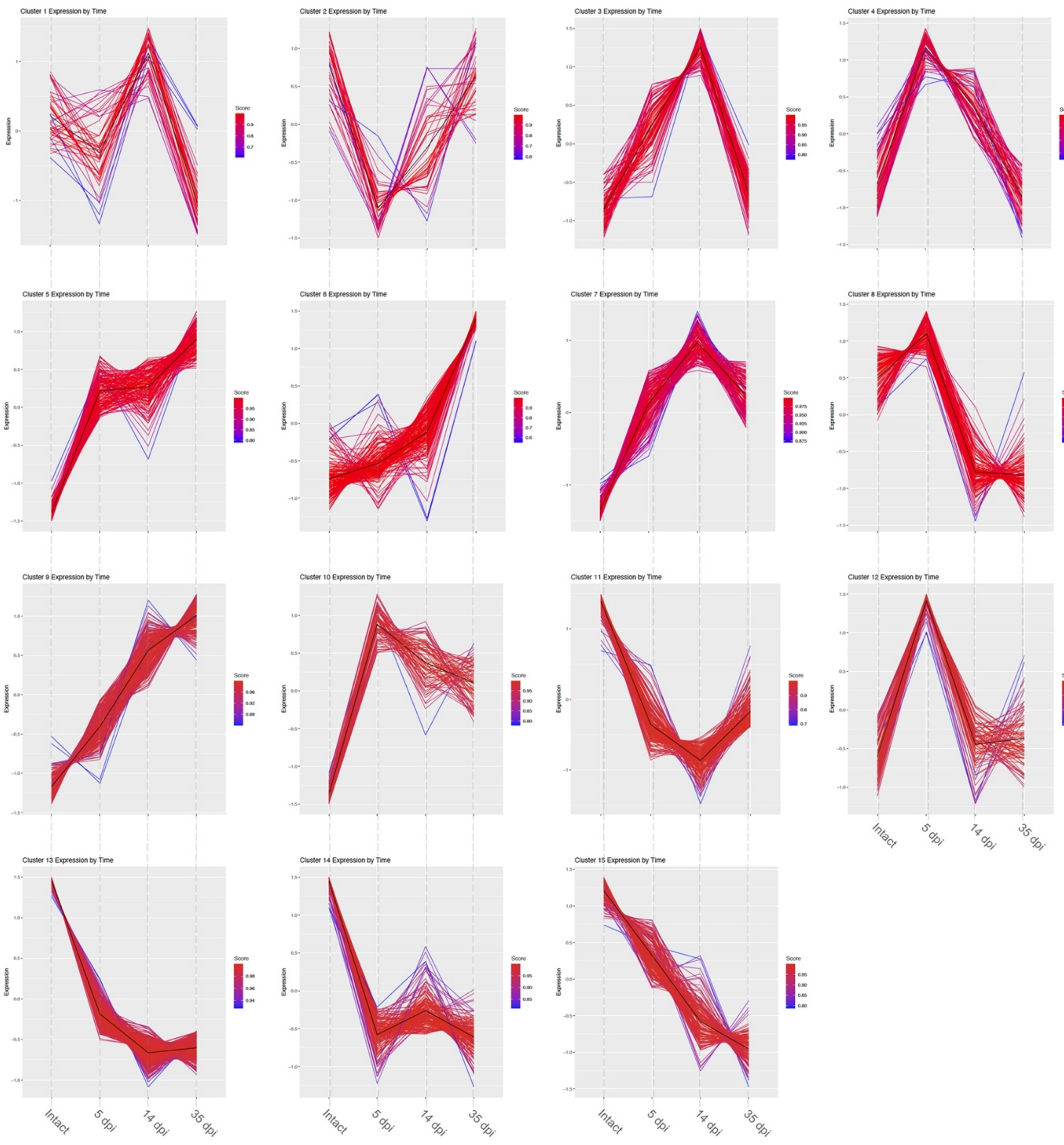


**Fig. S3 15 clusters capturing the gene expression of lower jaw regeneration in intact tissues and at 5-, 14-, and 35 dpi** Color gradient indicates the Pearson Correlation of any given gene to the core of its corresponding cluster. Y-axis indicates expression as scaled z-scores.

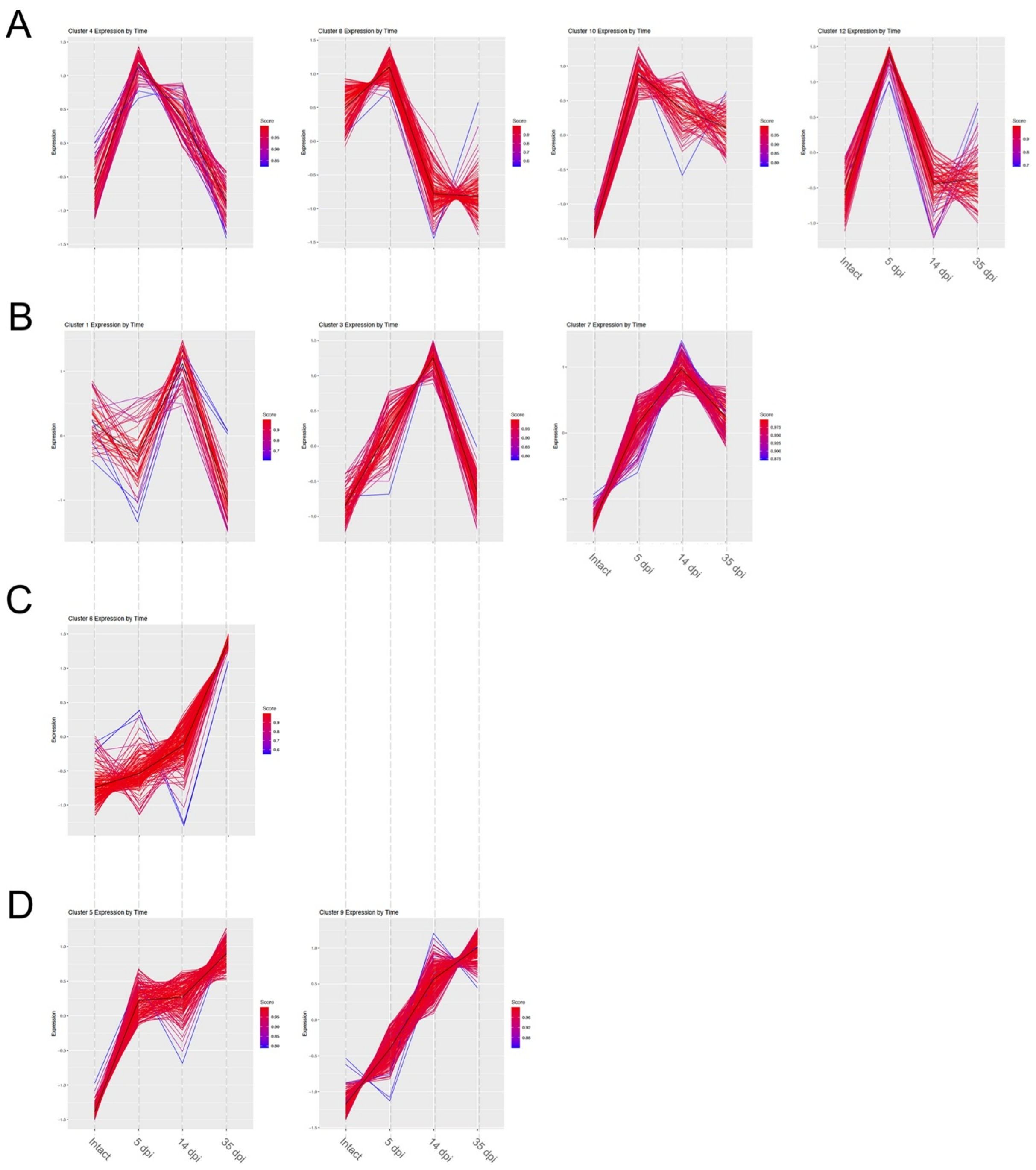


**Fig. S4 Regenerating lower jaw cluster groupings for downstream analysis. A.** Clusters 4, 8, 10 and 12 are grouped in the category *5 dpi Peak*. **B.** Clusters 1, 3, and 7 are grouped in the category *14 dpi Peak*. **C.** Cluster 9 is the only one composing the category *35 dpi Peak*. **D.** Clusters 6, and 9 are grouped in the category *General Rise*. Gradient indicates the Pearson Correlation of any given gene to the core of its corresponding cluster. Y-axis indicates expression as scaled z-scores.

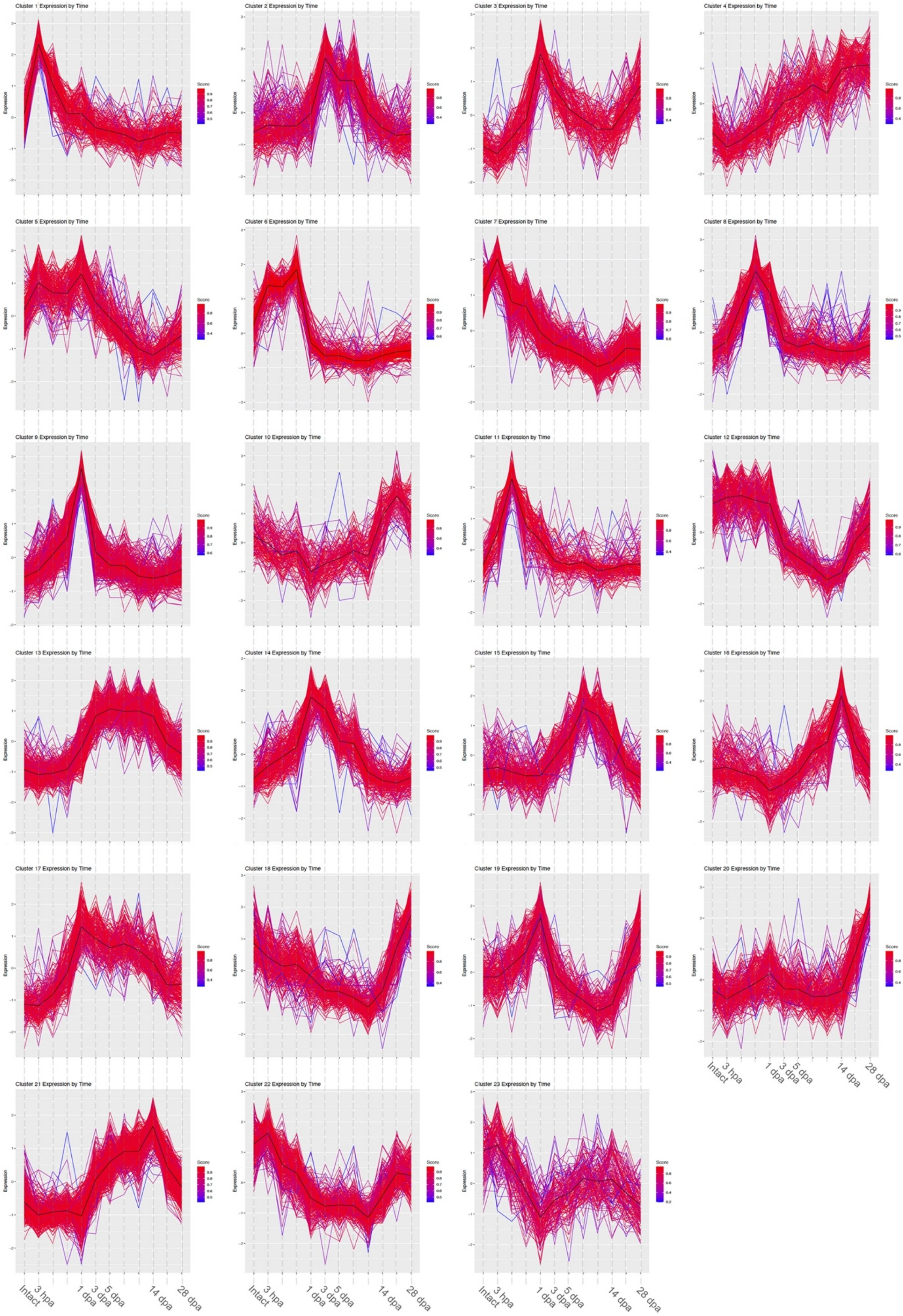
Cluster 1 Expression by Time
Cluster 2 Expression by Time
Cluster 3 Expression by Time
Cluster 4 Expression by Time
Cluster 5 Expression by Time
Cluster 6 Expression by Time
Cluster 7 Expression by Time
Cluster 8 Expression by Time
Cluster 9 Expression by Time
Cluster 10 Expression by Time
Cluster 11 Expression by Time
Cluster 12 Expression by Time
Cluster 13 Expression by Time
Cluster 14 Expression by Time
Cluster 15 Expression by Time
Cluster 16 Expression by Time
Cluster 17 Expression by Time
Cluster 18 Expression by Time
Cluster 19 Expression by Time
Cluster 20 Expression by Time
Cluster 21 Expression by Time
Cluster 22 Expression by Time
Cluster 23 Expression by Time
Expression
Score
Intact
3 hpa
1 dpa
3 dpa
5 dpa
14 dpa
28 dpa

**Fig. S5 23 clusters capturing the gene expression of limb regeneration over time found by *k*-means clustering algorithm.** Color gradient indicates the Pearson Correlation of any given gene to the core of its corresponding cluster. Y-axis indicates expression as scaled z-scores. Data from Stewart et al., 2013.

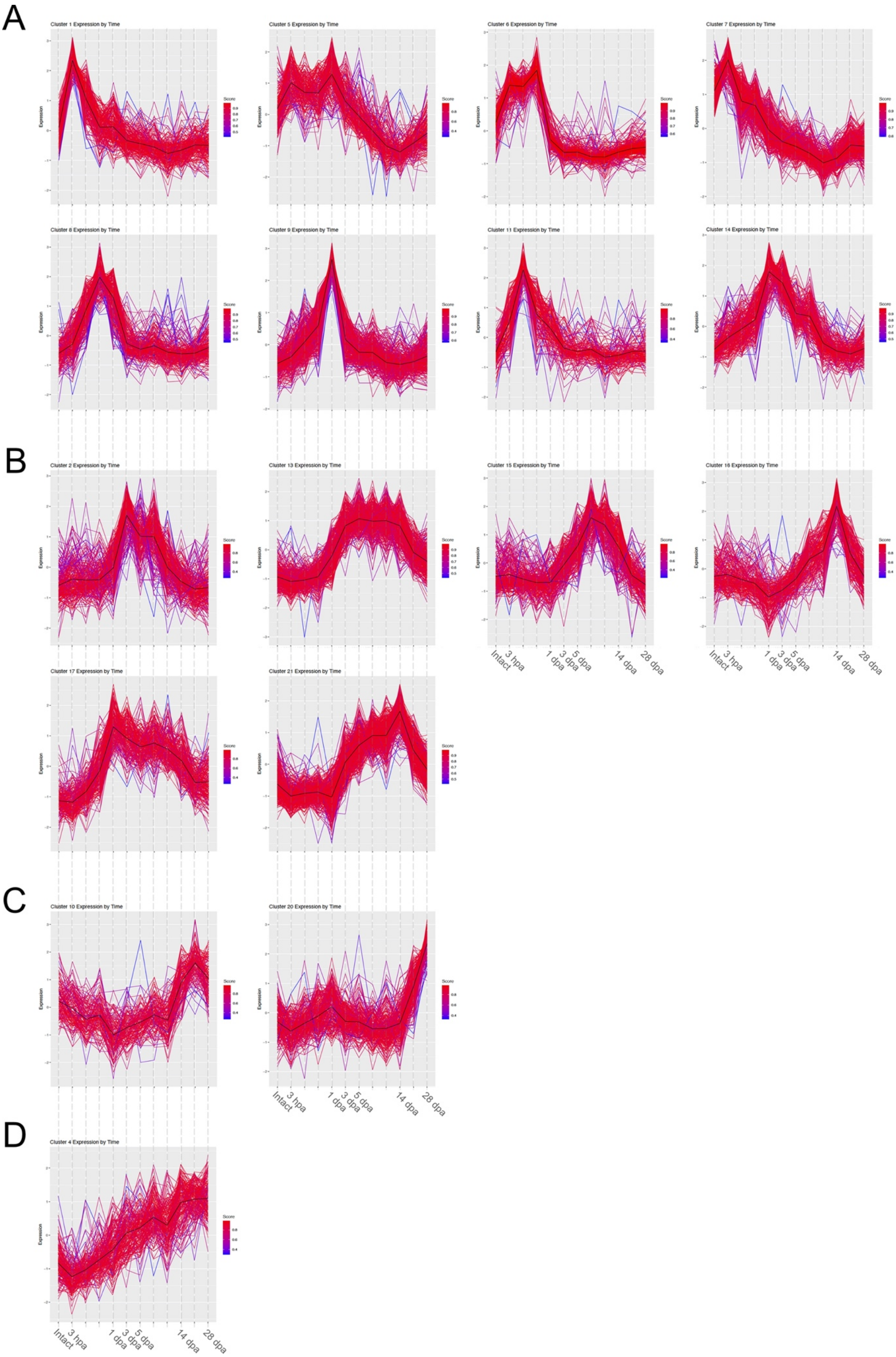
A
Cluster 1 Expression by Time
Cluster 5 Expression by Time
Cluster 6 Expression by Time
Cluster 7 Expression by Time
Cluster 8 Expression by Time
Cluster 9 Expression by Time
Cluster 11 Expression by Time
Cluster 14 Expression by Time
B
Cluster 2 Expression by Time
Cluster 13 Expression by Time
Cluster 15 Expression by Time
Cluster 16 Expression by Time
Cluster 17 Expression by Time
Cluster 21 Expression by Time
C
Cluster 10 Expression by Time
Cluster 20 Expression by Time
D
Cluster 4 Expression by Time
Expression
Score
Intact
3 hpa
1 dpa
3 dpa
5 dpa
14 dpa
28 dpa

**Fig. S6 Regenerating limb cluster groupings for downstream analysis. A.** Clusters 1, 5, 6, 7, 8, 9, 11 and 14 are grouped in the category *Early Peak*. **B.** Clusters 2, 13, 15, 16, 17 and 21 are grouped in the category *Mid Peak*. **C.** Clusters 10 and 20 are grouped in the category *Late Rise*. **D.** Clusters 6 is the only one in the category *General Rise*. Gradient indicates the Pearson Correlation of any given gene to the core of its corresponding cluster. Y-axis indicates expression as scaled z-scores.
